# Dual delivery of nucleic acids and PEGylated-bisphosphonates *via* calcium phosphate nanoparticles

**DOI:** 10.1101/621102

**Authors:** Sofia Bisso, Simona Mura, Bastien Castagner, Patrick Couvreur, Jean-Christophe Leroux

## Abstract

Despite many years of research and a few success stories with gene therapeutics, efficient and safe DNA delivery remains a major bottleneck for the clinical translation of gene-based therapies. Gene transfection with calcium phosphate (CaP) nanoparticles brings the advantages of low toxicity, high DNA entrapment efficiency and good endosomal escape properties. The macroscale aggregation of CaP nanoparticles can be easily prevented through surface coating with bisphosphonate conjugates. Bisphosphonates, such as alendronate, recently showed promising anticancer effects. However, their poor cellular permeability and preferential bone accumulation hamper their full application in chemotherapy. Here, we investigated the dual delivery of plasmid DNA and alendronate using CaP nanoparticles, with the goal to facilitate cellular internalization of both compounds and potentially achieve a combined pharmacological effect on the same or different cell lines. A pH-sensitive poly(ethylene glycol)-alendronate conjugate was synthetized and used to formulate stable plasmid DNA-loaded CaP nanoparticles. These particles displayed good transfection efficiency in cancer cells and a strong cytotoxic effect on macrophages. The *in vivo* transfection efficiency, however, remained low, calling for an improvement of the system, possibly with respect to the extent of particle uptake and their physical stability.

**Graphical abstract:** 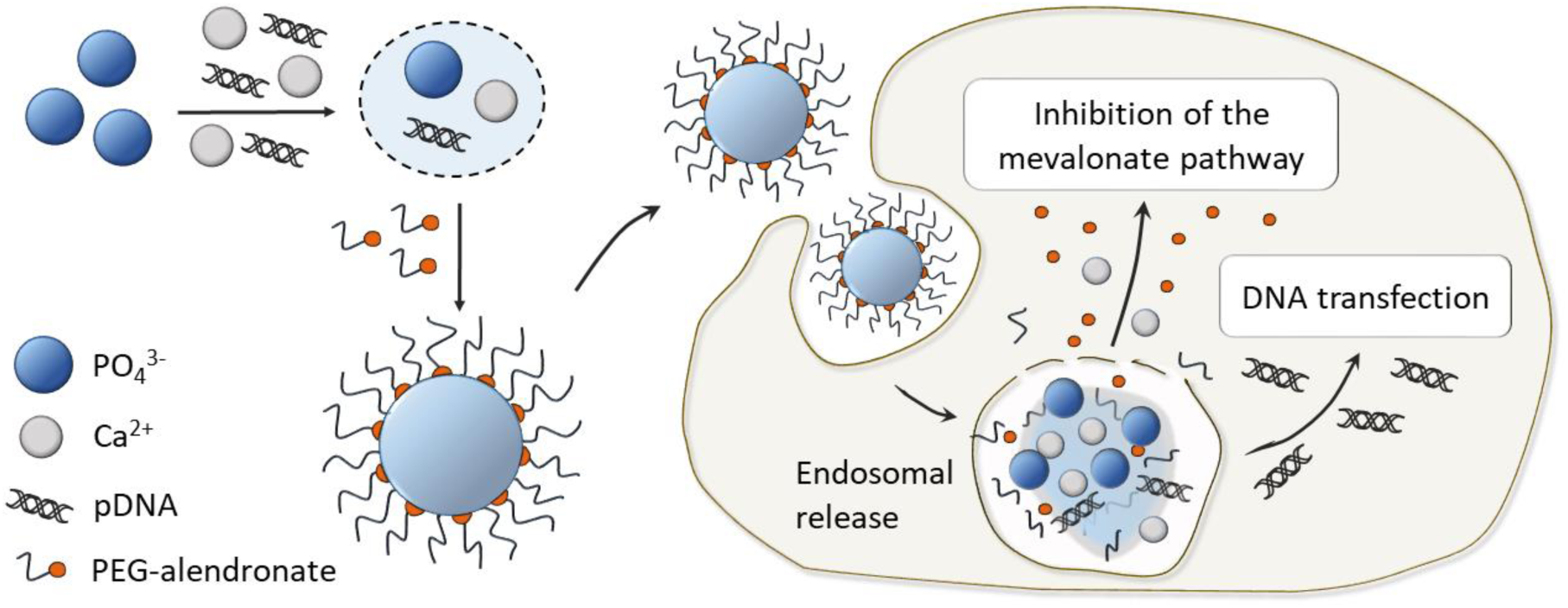

## 1. Introduction

Gene therapy represents a promising therapeutic modality for the treatment of inherited genetic disorders and other severe acquired diseases, such as cancer [1]. Despite their huge therapeutic potential, the clinical translation of nucleic acids is in part hampered by their high molecular weight, their polyionic nature and the susceptibility to degradation by nucleases, which restrict their biodistribution and permeability across cell membranes [2]. To overcome the hurdles associated to gene delivery, extensive research efforts have been put in the development of particle-based non-viral carriers, owing to their ability to shield DNA molecules from degradation and facilitate cellular uptake [3,4]. The ideal nanosized vehicle should selectively and efficiently deliver a gene to target cells with minimal toxicity. Additional desirable properties are ease of production, long shelf-life, high payload for genetic material and versatility of surface functionalization [4–7]. The most investigated and successful physicochemical methods for DNA delivery involve its electrostatic complexation to polycations (polyplexes) and/or cationic lipids (lipoplexes) [8]. However, these systems often display toxicity issues (necessitating premedication) and, unless sterically stabilized, they aggregate in physiological fluids leading to inefficient gene delivery at target sites [9]. Non-ionic particulate vehicles, based on lipids [10,11], biodegradable polymers [3,12] or inorganic nanoparticles (NPs) [3,5,13] have been described as alternative strategies to potentially improve gene delivery, safety and efficiency. Among them, calcium phosphate (CaP) NPs are well-established for the *in vitro* delivery of genetic material [14,15]. CaP-mediated transfection is based on the electrostatic entrapment of nucleic acid molecules in a relatively non-toxic and dissolvable particulate system, which eventually enhances the endocytosis and endosomal escape of the gene [16]. Nevertheless, rapid aggregation of CaP NPs upon preparation has always represented a significant hurdle for subsequent *in vivo* applications. Different methods have been investigated to prevent particle growth and agglomeration, mostly involving surface electrostatic stabilization with PEGylated polyanions [17], lipids [18], DNA/siRNA [19] and bisphosphonates (BPs) [16,20,21]. Nitrogen-containing BPs, such as alendronate (Ale), possess high affinity for calcium-containing matrices, and are conventionally used for the treatment of bone-related diseases [22,23]. Recently, BPs have attracted much attention for their direct and indirect anticancer benefits exerted on several non-skeletal tumours, such as breast and prostate cancer [24,25]. Among their multiple effects on the tumour microenvironment (i.e. on tumour associated macrophages), BPs can suppress cell proliferation *via* induction of apoptosis, primarily through the blockade of the mevalonate pathway [25,26]. Their action is mainly due to inhibition of the enzyme farnesyl pyrophosphate synthase (FPPS), which results in decreased prenylation of small guanosine triphosphate (GTP)-binding proteins (GTPases), ultimately leading to dysregulation of cell-signalling pathways [26,27]. However, the use of BPs as anticancer drugs still encounters limitations related to their rapid distribution to bones and poor membrane permeability. The dual-delivery of Ale and nucleic acids *via* a particulate system represents a potential strategy to increase the intracellular concentration of both therapeutic agents at target sites. Furthermore, by addressing two different intracellular pathways, the system may elicit a combined effect on therapeutic targets (e.g. cancer cells and/or tumour associated macrophages). For this purpose, a carrier with high adsorptive capacity for functional charged molecules must be designed. We previously reported that nucleic acid-loaded CaP NPs coated with a non-cleavable polyethylene glycol (PEG)*-*Ale derivative induced a modest increase in the intracellular levels of two toxic metabolites, isopentanylpyrophosphate (IPP) and triphosphoric acid 1-adenosin-5’-yl ester 3-(3-methylbut-3-enyl) ester (ApppI) [20]. Accumulation of these molecules is a consequence of inhibition of FFPS, and the effect was attributed to a putative partial enzymatic cleavage of PEG-Ale. Hence, we hypothesized that conjugation of Ale to PEG through an acid-cleavable linker could allow for a more quantitative release of the BP inside the cell upon endosomal acidification, thus potentially triggering cell death. In this work, we report a CaP-based nanoparticulate carrier for the delivery of both a BP drug and plasmid DNA to cells. A pH-sensitive PEG-Ale chelator was developed to stabilize the NPs, and Ale was conjugated to PEG *via* a maleic anhydride linker, which is hydrolyzed in mildly acidic conditions. The cellular uptake and transfection efficiency of CaP NPs were evaluated *in vitro,* and their ability to interfere with the mevalonate pathway was investigated both in macrophages and cancer cells. Finally, the *in vivo* transfection efficacy of CaP NPs was tested in mice harbouring orthotopic mammary tumours after local administration.

## 2. Materials and Methods

### 2.1 Materials

Ale sodium trihydrate and tris[(1-benzyl-1H-1,2,3-triazol-4-yl)methyl]amine (TBTA) were purchased from Tokyo Chemical Industry Co., Ltd. (Tokyo, Japan). Cobalt thiocyanate was obtained from Siegfried (Zofingen, Switzerland). Triethylamine and 5-hehyn-1-ol 97% were purchased from Acros Organics (Geel, Belgium). Deuterium oxide (D_2_O) and deuterated chloroform were obtained from Cambridge Isotope Laboratories (Tewksbury, MA). Methoxy-PEG succinimidyl carboxymethyl ester (M_n_ ≈ 2000 Da) was bought from JenKem Technology USA (Plano, TX). Sephadex G-25 PD-10 desalting columns were from GE Healthcare life science (Chalfont St. Giles, UK). 4T1 murine breast cancer cells and J774.2 murine macrophages were purchased from ATCC (Manassas, VA). Dulbecco’s modified Eagle medium (DMEM) with GlutaMAX™ (high glucose), phenol red-free DMEM, Opti-MEM™ I reduced serum medium, fetal bovine serum (FBS), penicillin/streptomycin stock solution (10,000 IU/mL penicillin and 10,000 µg/mL streptomycin), trypsin 0.25 % (*m/v*)/1 mM EDTA solution, phosphate buffered saline (PBS), Live Cell Imaging Solution for cell culture, Hoechst 33342 and ProLong™ Diamond antifade mountant were obtained from Life Technologies (ThermoFisher Scientific, Carlsbad, CA). MycoAlert PLUS Mycoplasma Detection Kit was purchased from Lonza (Basel, Switzerland). *Label* IT^®^ plasmid delivery control, fluorescein labeled pDNA (2.7 kb), was obtained from LabForce (Muttenz, Switzerland). X-tremeGENE™ 9 DNA Transfection Reagent was bought from Roche Pharma (Reinach, Switzerland). CellTiter 96 AQueous One Non-Radiactive Cell Proliferation Assay based on 3-(4,5-dimethylthiazol-2-yl)-5-(3-carboxymethoxyphenyl)-2-(4-sulfophenyl)-2H-tetrazolium (MTS) was from Promega (Duebendorf, Switzerland). COmplete™ EDTA-free protease inhibitor cocktail was obtained from Roche Diagnostics (Mannheim, Germany). Immun-Blot^®^ poly(vinylidene difluoride) membranes were purchased from Bio-Rad Laboratories (Hercules, CA). Mouse monoclonal Rap1A (C-10) antibody, mouse monoclonal β-actin (C4) antibody and ImmunoCruz^®^ Western blotting Luminol Reagent were obtained from Santa Cruz Biotechnology (Dallas, TX). Horseradish peroxidase (HRP)-conjugated goat anti-mouse IgG polyclonal antibody was purchased from Dako (Glostrup, Denmark). Super RX X-ray films were obtained from Fujifilm (Tokyo, Japan). *In vivo*-jetPEI^®^ was purchased from Polyplus-transfection S.A (Illkirch, France). Roti^®^ Histofix 4% PFA solution was from Carl Roth (Karlsruhe, Germany) and optimal cutting temperature (OCT) embedding medium was bought from CellPath (Powys, United Kingdom). ATTO488-GFP-Booster was purchased from ChromoTek (Planegg-Martinsried, Germany). All other chemicals were obtained from Sigma-Aldrich (Buchs, Switzerland).

### 2.2 Synthesis and characterization of PEG-Ale coating agent

Synthesis of pH-sentitive PEG-2-(pent-1′-yne) maleic acid Ale amide (PEG-ma-Ale), PEG-1-methyl-2-(2′-carboxyethyl) maleic acid Ale amide (PEG-mma-Ale) and the pH-stable PEG-amide-Ale (PEG-a-Ale) control are described in detail in the supplementary information section (Fig. **S1-S10**).

### 2.3 Determination of Ale release from PEG-Ale prodrugs

The kinetics of Ale release from the PEG-ma-Ale and PEG-mma-Ale maleic acid amide prodrugs were monitored at room temperature (RT) by ^1^H-NMR spectroscopy at different pH values. Each derivative was dissolved at a 20 mg/mL concentration in D_2_O-based 20 mM Na_2_HPO_3_ buffer, and the pH of the solution was adjusted to 7.4 or 5.0 with 1N NaOD or 1N DCl. For the measurement at basic pH values, a 20-mM Na_2_CO_3_ buffer was used, and the pH was adjusted to 9.0 by addition of 1N NaOD or 1N DCl. For the hydrolysis experiments on PEG-ma-Ale performed at pH 5.0, an aliquot of the prodrug was incubated in acidic buffer for each time point, followed by purification *via* size-exclusion chromatography using a Sephadex G-25 PD-10 column (eluent 20 mM Na_2_HPO_3_ buffer, pH 8.5). The purified PEG-ma-Ale derivative was lyophilized and subsequently resuspended at a 20 mg/mL concentration in D_2_O-based 20 mM Na_2_HPO_3_, pH 7.4 for NMR analysis of hydrolysis. For each time point, semi-quantitative degradation analysis on PEG-ma-Ale was calculated according to Eq. (1):

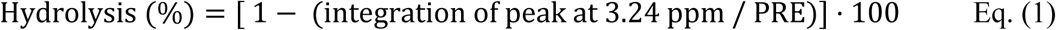

where PRE is the value of integration of peak at 3.24 ppm at time 10 min of PEG-ma-Ale dissolved in D_2_O-based 20 mM Na_2_HPO_3_ buffer, pH 7.4 (Fig. **S11**). For comparison purposes between NMR spectra, integration of the peak at 2.6 ppm was normalized to 2 H for each time point.

In the same way, the extent of hydrolysis of PEG-mma-Ale was calculated according to Eq. (2)>:

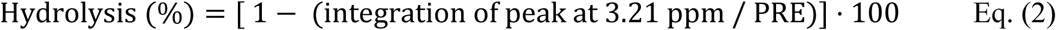

where PRE is the value of integration of peak at 3.21 ppm at time 10 min of PEG-mma-Ale dissolved in D_2_O-based 20 mM Na_2_CO_3_ buffer, pH 10 (Fig. **S14**). To compare NMR spectra, integration of the peak at 4.29 ppm was normalized to 2 H for each time point.

### 2.4 Plasmid DNA amplification

Plasmid DNA (pc3DNA; enhanced green fluorescent protein (EGFP) insert, ampicillin resistance) from Addgene (Cambridge, MA) was amplified in DH5α E. coli and then extracted and purified using a plasmid purification maxi kit (Qiagen, Valencia, CA). Purity was assessed by the ratio of absorbance at 260 and 280 nm, using a NanoPhotometer spectrophotometer (Implen, München, Germany). Plasmids with an absorbance ratio of ∼1.8 were considered as pure.

### 2.5. Particle preparation

CaP NPs were prepared according to reference [21]. Briefly, one volume of 50 mM HEPES buffer (containing 140 mM NaCl and 1.5 mM Na_2_HPO_3_, pH 7.4) was mixed with one volume of a 250-mM CaCl_2_ solution containing 20 µg/mL pEGFP, under vigorous agitation. After particles nucleation, one volume of 10 or 20 µM PEGylated chelator in 10 mM TRIS, pH 7.4, was quickly added to stabilize the particles. For cytotoxicity assays, additional solutions with 40, 50 and 80 µM of PEGylated chelators were used. CaP NPs for *in vivo* studies were prepared according to the standard method using buffers with a lower ionic strength (*vide infra*) to limit the increase in osmolality after particles concentration [28]. One volume of 7 mM HEPES buffer (containing 18 mM NaCl and 1.5 mM Na_2_HPO_3_, pH 7.4) was mixed with one volume of 125 mM CaCl_2_ (containing 40 µg/mL pEGFP), followed by quick addition of one volume of 15 µM PEG-ma-Ale in 2 mM TRIS, pH 7.4. Subsequently, 375 µL of NP suspension was mixed 1:1 (*v/v*) with a 26 mg/mL trehalose solution in ultrapure water, snap-frozen in liquid nitrogen and lyophilized. For the *in vivo* experiments, the freeze-dried CaP NPs were reconstituted in ultrapure water to a final volume of 50 µL, vortexed shortly and sonicated for 5 min.

### 2.6. Particle characterization

Particle size, polydispersity index (PDI) and ζ-potential were determined by dynamic light scattering (DLS) and laser Doppler anemometry respectively, using a DelsaNano C Particle Analyzer (Beckman Coulter, Krefeld, Germany). The CONTIN method was used to calculate the hydrodynamic diameter of the particles. ζ-potential was measured after dilution of the particles 1:2 (*v/v*) in 10 mM TRIS, pH 7.4. Osmolality was measured with a freezing point osmometer (Osmomat 3000, Gonotec GmbH, Berlin, Germany). Transmission electron microscopy (TEM) of the particle suspension was performed at 100 kV (FEI Morgagni268, FEI Company, Hillsboro, OR). To calculate the encapsulation efficiency, *Label* IT^®^ plasmid fluorescein delivery control was loaded on CaP NPs (225 µL, final pDNA concentration of 6.66 µg/mL). The particles were then isolated by centrifugation (30 min, 4 °C, 18,000 *x g*) and 180 µL of supernatant were taken to measure the fluorescence intensity of free pDNA (λ_ex_ = 485 nm, λ_em_ = 520 nm) with a plate reader (Infinite M-200, Tecan, Männedorf, Switzerland). Encapsulation efficiency (EE) was estimated by subtracting the free DNA concentration in the supernatant from the total concentration, using the fluorescence signals (FI), according to Eq. (3):

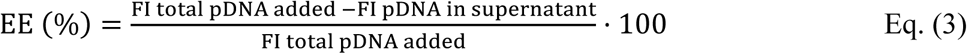

### 2.7 Colloidal stability of CaP NPs

Stability of 10 µM PEG-ma-Ale-coated CaP NPs at different pHs was evaluated over time by DLS after adjusting the pH to 7.0, 6.8 or 6.5 by addition of 1N NaOH or 1N HCl. The stability in serum-containing media of pEGFP loaded NPs (pDNA concentration: 6.66 µg/mL) prepared with 10 and 20 µM feeding amount of PEG-ma-ALE was assessed overtime at 37 °C in complete medium containing FBS. CaP NPs were diluted 1:7 (*v/v*) in DMEM containing 10 or 50% FBS, and their hydrodynamic diameter was recorded over time by DLS at 37 °C. Additionally, the stability of pEGFP-loaded CaP NPs coated with different PEG-ma-Ale feeding amounts (10, 15 and 20 µM) was measured by DLS at RT at defined time intervals, for up to 21 days.

### 2.8. Cell culture

4T1 (murine mammary carcinoma) and J774.2 (murine macrophage-like) cells were grown in DMEM with GlutaMAX™ (high glucose) supplemented with 10% (*v*/*v*) FBS, 100 IU/mL penicillin and 100 µg/mL streptomycin. Cells were kept at 37 °C in a humidified atmosphere containing 5% CO_2_. Cells were used from passages 3 to 20, and regularly checked for mycoplasma contamination.

### 2.9. *In vitro* particles uptake

The uptake of pDNA loaded CaP NPs stabilized with PEG-ma-Ale was evaluated in 4T1 and J774.2 cells. In order to track the internalized particles, CaP NPs were loaded with *Label* IT^®^ Fluorescein Plasmid Delivery Control as model pDNA. 4T1 and J774.2 cells were seeded into 24-well tissue-treated plates at a density of 60,000 and 100,000 cells/well, respectively, in 0.5 mL complete DMEM containing 10% FBS. After 24 h incubation at 37 °C in a humidified atmosphere containing 5% CO_2_, the medium was changed to complete DMEM supplemented with 10% FBS, and particles prepared with 10 or 20 µM feeding amounts of PEG-ma-ALE were diluted 1:6.66 (*v/v*) in complete medium to a final pDNA concentration of 1 µg/mL. After 5 h or 24 h incubation, the medium was removed, and cells were washed twice with 200 µL PBS. Then, 4T1 cells were detached by trypsinization, followed by addition of complete medium to stop the trypsin action. To detach J774.2 cells, 0.5 mL of complete medium were added to each well, and cells were recovered with a cell scraper. Cells were isolated by centrifugation (10 min, 4 °C, 300 *x g*) and subsequently resuspended in ice cold FACS buffer (PBS, 2 mM EDTA, 0.5% *m/v* BSA). Data for 10,000 cells were collected on a CytoFLEX Flow Cytometer (Beckman Coulter Inc., Brea, CA) and analyzed using the FlowJo software (Tree Star Inc., Ashland, OR). Uptake of CaP NPs was determined as percentage of FITC-positive cells, compared to control wells that did not receive CaP NPs. For the uptake in the inverted configuration, 4T1 and J774.2 cells were seeded on 13-mm Nunc™ Thermanox™ Coverslips (ThermoFisher Scientific, Carlsbad, CA) in a 24-well plate at a density of 60,000 and 100,000 cells/well respectively. After 24 h incubation at 37 °C in a humidified atmosphere containing 5% CO_2_, 0.5 mL of complete medium containing 10% FBS were added to each well of a 24-well plate, and CaP NPs containing fluorescein-labeled pDNA were added at a 1:6.66 (*v/v*) final dilution, corresponding to a final pDNA concentration of 1 µg/mL. The coverslips on which cells adhered were carefully transferred upside down onto the surface of the medium. After 24 h, the medium was removed and the coverslips were transferred in a 24-well plate and washed twice with PBS. Cells were prepared for FACS analysis and data were processed as previously described for the uptake in upright configuration.

### 2.10. *In vitro* particles transfection efficiency

4T1 and J774.2 cells were seeded into 24-well tissue-treated plates at a density of 20,000 and 80,000 cells/well, respectively, in 0.5 mL complete DMEM containing 10% FBS. After 24 h incubation at 37 °C in humidified atmosphere containing 5% CO_2_, the medium was removed and the cells were incubated with CaP NPs formulations diluted 1:6.66 (*v/v*) in Opti-MEM containing 10% FBS, to a final pDNA concentration of 1 µg/mL. Cells were exposed to PEG-ma-Ale stabilized CaP NPs for different incubation times (5, 24 or 48 h). X-tremeGENE™ was used as positive control, as well as not stabilized CaP NPs. After the 5 and 24 h incubation time points, the medium containing the CaP NPs was removed, the cell layer was washed twice with PBS and then incubated with 500 µL complete medium for up to 48 h after particles addition. Then, the medium was aspirated and the cell layer was washed twice with PBS. Cells were prepared for FACS analysis and data were processed as described in section 2.9. EGFP expression was evaluated as the percentage of FITC-positive cells, compared to control wells not exposed to CaP NPs. Fluorescence images of EGFP expressing cells were acquired with a LEICA DMI600B epifluorescence microscope (Leica Microsystem, Wetzlar, Germany).

### 2.11. *In vitro* particles cytotoxicity

The viability of 4T1 and J774.2 cells after incubation with CaP NPs stabilized with the different PEG-Ale stabilizing agents was evaluated by the MTS assay. 4T1 and J774.2 cells were seeded into flat-bottomed 96-well plates at a density of 5000 and 10,000 cells/well, respectively, in 100 µL complete DMEM containing 10% FBS. After 24 h incubation, the medium was removed and the cells were incubated with different formulations of CaP NPs diluted 1:6.66 (*v/v*) in complete medium, to achieve different final concentrations of each PEG-ma-Ale or PEG-a-Ale stabilizing agent (0.5, 1, 2, 4, and 5 µM, calculated based on the corresponding Ale molar concentrations). Cells were exposed to the CaP NPs for different incubation times (5, 24 or 48 h). As control, cells were incubated with only PEG-ma-Ale. After the 5 and 24 h incubation time points, the medium containing the CaP NPs was removed, the cell layer was washed twice with PBS, and 100 µL of complete medium were added to each well for up to 48 h from particles addition. Then, the medium was aspirated, the cell layer was washed twice with PBS and 120 µL of DMEM without phenol red containing the MTS reagent in the ratio 6:1 (*v/v*) were added. After 1 h incubation at 37 °C, the absorbance of the wells was measured at 490 nm with a Tecan plate reader. Cell viability was determined according to Eq. (4):

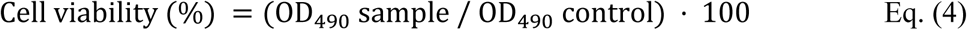

where OD_490_ sample represents the optical density (OD) of the wells treated with CaP NPs, and OD_490_ control represents the wells treated with growth medium only. One mM hydrogen peroxide was used as positive toxic control (viability < 20%), and wells containing only reaction mixture but no cells were used as blank.

### 2.12. Western Blot analysis of Rap1A

4T1 and J774.2 cells were seeded in 12-well plates at 40,000 and 80,000 cells/well, respectively, in 1 mL complete DMEM containing 10% FBS. Twenty-four hours after seeding, CaP NPs stabilized with PEG-ma-Ale or PEG-a-Ale were added to each well at a 1:6.66 (*v/v*) final dilution, corresponding to different PEG-Ale concentrations (0.5, 2 and 4 µM, calculated based on the corresponding Ale concentrations). Complete medium without NPs was used as negative control. After 24 h incubation, the medium was removed, cells were washed twice with PBS and cell lysates were prepared in 120 µL lysis buffer (20 mM Tris–HCl pH 7.7, 150 mM NaCl, 5 mM EDTA, 1% *v/v* Triton X-100, 25 mM NaF, 1 mM phenylmethylsulfonyl fluoride, 1 mM sodium orthovanadate), supplemented with cOmplete™ EDTA-free protease inhibitor cocktail, on ice for 15 min. Cell debris were removed by centrifugation (15 min, 4 °C, 15,000 × *g*) and protein concentration in the supernatants was determined using the BCA assay (ThermoFisher Scientific, Waltham, MA). Ten micrograms of total protein per sample were resolved by a 12% SDS-PAGE and transferred to a PVDF membrane. The membrane was blocked with Tris-buffered saline containing 0.1% (*v/v*) polysorbate 20 and 5% (*m/v*) skim milk (blocking buffer) for 1 h at RT. The membrane was cut in two at 35 kDa, determined by molecular weight markers. The lower part containing Rap1A (22 kDa) was incubated overnight at 4 °C with mouse Rap1A monoclonal antibody diluted 1:500 in blocking buffer. The upper part containing β-actin (internal control) was incubated with mouse polyclonal anti-β-actin antibody at a dilution of 1:500 in blocking buffer. Membranes were washed with Tris-buffered saline containing 0.1% (*v/v*) polysorbate 20, and then incubated for 2 h with the secondary antibody, horseradish peroxidase-conjugated polyclonal goat anti-mouse IgG, diluted 1:1000 in blocking buffer. Membranes were washed with Tris-buffered saline containing 0.1% (*v/v*) polysorbate 20 and protein bands were detected with ImmunoCruz^®^ Western blotting Luminol Reagent and revealed on Super RX X-ray films, using an AGFA Curix 60 film processor (AGFA, Mortsel, Belgium).

### 2.13 *In vivo* transfection efficiency study

All animal experiments were approved by the Animal Care Committee of the University Paris-Sud (registered under no. 11894/2017102312072785). Experimental procedures were performed according to the 2010/63/EU European directive and the instructions of laboratory animal care and legislation in force in France. Female BALB/c mice (6-8 weeks) were purchased from Janvier Labs (Le Genest-Saint-Isle, France). To establish the orthotopic mammary tumours, 4T1 cells were harvested using trypsin, isolated by centrifugation at 200 x *g* x 5 min and resuspended in PBS at a density of 1×10^6^ cells/mL. Mice were anesthetized under isoflurane (3% *v/v* in 1 mL/min oxygen flow) and 50 µL of the cell suspension were injected in the fourth inguinal mammary fat pad, using a 29-G syringe (Terumo, Tokyo, Japan). Tumour size was regularly measured with a caliper, and its volume was estimated using Eq. (5):

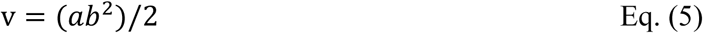

where *a* is the larger tumour dimension and *b* the smallest. When the tumours reached a volume of approximatively 100 mm^3^, mice were anaesthetized under isoflurane (3% *v/v* in 1 mL/min oxygen flow) and divided into 4 random groups of 3-8 animals, that received the following treatments: (i) pEGFP loaded CaP NPs, (ii) pEGFP in 5% (*m/v*) glucose, (iii) pEGFP complexed with *in vivo*-jetPEI^®^ with a N/P ratio of 6, in 5% (*m/v*) glucose (positive control), (iv) 5% (*m/v*) glucose (negative control, n = 3). The formulations were slowly administered intratumorally in multiple spots, at a final pDNA dose of 2 µg in 20 µL. The mice received a total of three doses of 2 µg pDNA each, every other day over one week. After 24 h from the last dose, mice were anaesthetized under isoflurane (3% *v/v* in 1 mL/min oxygen flow) and sacrificed *via* cervical dislocation. Excised tumour tissues were fixed in 4% PFA at 4 °C overnight, then washed twice with PBS and placed in 15% (*m/v*) sucrose in PBS at 4 °C until sinking. The specimens were then transferred in 30% (*m/v*) sucrose in PBS and stored at 4 °C overnight. The samples were embedded in OCT and subsequently frozen on a metal support placed in liquid nitrogen. The frozen samples were stored at −80 °C until cryosectioning procedure. Tissue cryosections (10 µm) were cut with a CryoStar NX70 (Thermo Fisher Scientific, Waltham, MA) and the slices were mounded on SuperFrost Plus glass slides (Thermo Fisher Scientific, Waltham, MA). The slices were fixed and permeabilized in ice cold methanol for 15 min, then blocked in 5% (*m/v*) BSA in PBS for 30 min. ATTO488-GFP-Booster was added at a 1:200 dilution in blocking buffer, followed by overnight incubation at 4 °C. The slices were stained for 20 min with Hoechst 33342 added at a 1:2000 dilution in PBS, then mounted on a coverslip using ProLong™ Diamond and visualized with a Leica epifluorescence microscope.

### 2.14. Statistical analysis

All values were expressed as mean ± standard deviation. All statistical analyses were performed using the GraphPad Prism software, version 8. Normality was evaluated with the Kolmogorov-Smirnov test. When data followed normal distribution, pairwise comparison between multiple groups was performed using the one-way analysis of variance (ANOVA) test, followed by Tukey’s (Holm-Sidak) post-hoc test. For non-normal distribution (e.g., cell viability assays) treatment groups were compared with one-way Kruskall–Wallis test followed by Dunn’s post-hoc test. Results were considered statistically significant at *p* ≤ 0.05.

## 3. Results and Discussion

### 3.1 Synthesis and characterization of the PEGylated chelators

The PEG-ma-Ale derivative was synthetized by conjugation of Ale to one PEG molecule, using a monosubstituted maleic anhydride as acid-cleavable linker. The cyclic anhydride was synthetized *via* a three-step procedure from a terminal alkyne bearing a primary alcohol, which was converted to aldehyde, followed by Mannich cyclization and oxidation to cyclic anhydride. Then, a maleic acid amide was obtained by coupling with Ale under basic conditions, and a PEG chain was introduced through a copper(I)-catalysed alkyne–azide cycloaddition reaction, to finally afford PEG-ma-Ale (Fig. 1a). In order to compare the Ale release rate from a different anhydride linker, a disubstituted maleic anhydride linker, *i.e.* 2-propionic-3-methylmaleic anhydride (here named mma), was coupled to one PEG molecule *via* an ester bond and then conjugated to Ale, to yield PEG-mma-Ale (Fig. 1b). The PEG-a-Ale stable derivative was obtained by conjugation of Ale and PEG *via* an amide bond, as previously reported (Fig. **S10**) [20]. The hydrolysis rate of the two pH-sensitive PEG-Ale derivatives at various pH values was measured over time by ^1^H-NMR spectroscopy. A negligible (< 10%) Ale release was observed after 24 h incubation of PEG-ma-Ale both at pH 9.0 and 7.4 (Fig. 2a and **S11**). The compound slowly hydrolyzed at pH 5.0 (Fig. **S12**), with ∼50% Ale released after about 3 h (Fig. 2a). The amide was almost fully cleaved (> 80% hydrolysis) within 1 h at pH 4.0 (Fig. 2a). At this pH value, Ale release was complete after 6 h incubation, as inferred from the disappearance of the methylene protons next to the nitrogen in the intact amide in the NMR spectrum (at 3.24 ppm) (Fig **S13**). Conversely, Ale was quickly released from PEG-mma-Ale at pH 7.4 (Fig. **S14**), with ∼50% of hydrolysis reached after about 50 min (Fig. 2b), while a slow hydrolysis was observed at pH 9.0, with 50% of the Ale released within 24 h. In cells, the pH varies significantly depending on the cellular compartment, ranging from 8 in mitochondria to as low as 5.0 to 4.0 in endosomes and lysosomes. Moreover, the pH is generally slightly lower in tumour mass (pH ≈ 6.5-7.2) than in blood and normal tissues [29]. The optimal pH sensitive moiety for intracellular release should be stable in biological fluids and cleavable within the cell, exploiting mild pH changes. Maleic anhydrides react with amines to yield maleic acid amides, which are cleaved in slightly acidic environments, when the nitrogen of the amide bond is in equilibrium with its protonated form. The whole cleavage process is driven by intramolecular catalysis of the anhydride that favors the formation of the cyclic form (Fig. **S15**) [30]. This peculiar characteristic of acid labile maleic anhydride derivatives was extensively exploited in a variety of pharmaceutical applications, such as to generate prodrugs for controlled drug release [31], or to design pH-dependent charge-conversion derivatives that confer endosomolytic properties to a carrier [32–34]. Depending on the nature of the substituents at the unsaturated carbon−carbon bond, as well as on the pK_a_ of the conjugated amine, the release rate of the amine can be finely tuned at acidic pH. For instance, Kang S. et al [35] observed that a faster amine release could be achieved at a higher degree of substitution at the *cis*-double bond of the anhydride ring, due to the Thorpe-Ingold effect [36,37]. Accordingly, the monosubstituted anhydride linker provided appropriate stability to PEG-ma-Ale at neutral pH, and controlled degradability at weak acidic pH. On the other hand, substitution of the hydrogen atom with a methyl group at the *cis*-double bond of PEG-mma-Ale accelerated the release rate of the BP at pH 7.4. Therefore, the conjugate obtained from the disubstituted anhydride was not selected for further studies, due to its predicted instability in formulation buffers and biological fluids.

**Figure 1.**
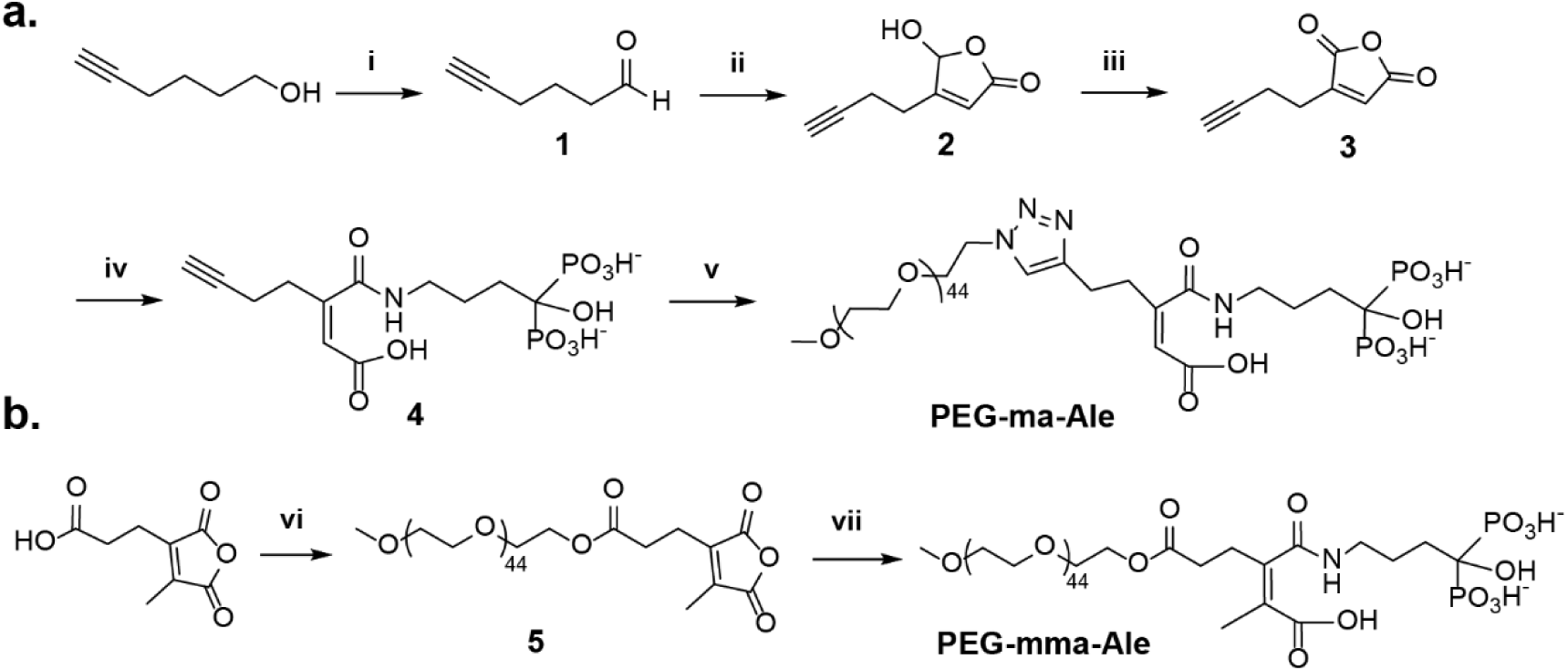
Synthesis of PEG-ma-Ale (a) and PEG-mma-Ale (b). Reagents and conditions: (i) 1. oxalyl chloride, dimethyl sulfoxide, dichloromethane (DCM), −78 °C; 2. triethylamine, −78 °C; (ii) Glyoxylic acid monohydrate, morpholine· hydrochloric acid salt, aq. dioxane, 100 °C, (iii) Dess-Martin periodinane, DCM, RT; (iv) Ale sodium trihydrate, water, pH 9; (v) mPEG-azide, copper(I) bromide, tris((1-benzyl-4-triazolyl)methyl)amine, aq. dimethyl sulfoxide/*tert*-butanol, pH 8.5; (vi) 1. oxalyl chloride, dimethylformamide, DCM; 2. mPEG-OH, DCM; (vii) Ale sodium trihydrate, water, pH 9-10.

**Figure 2.**
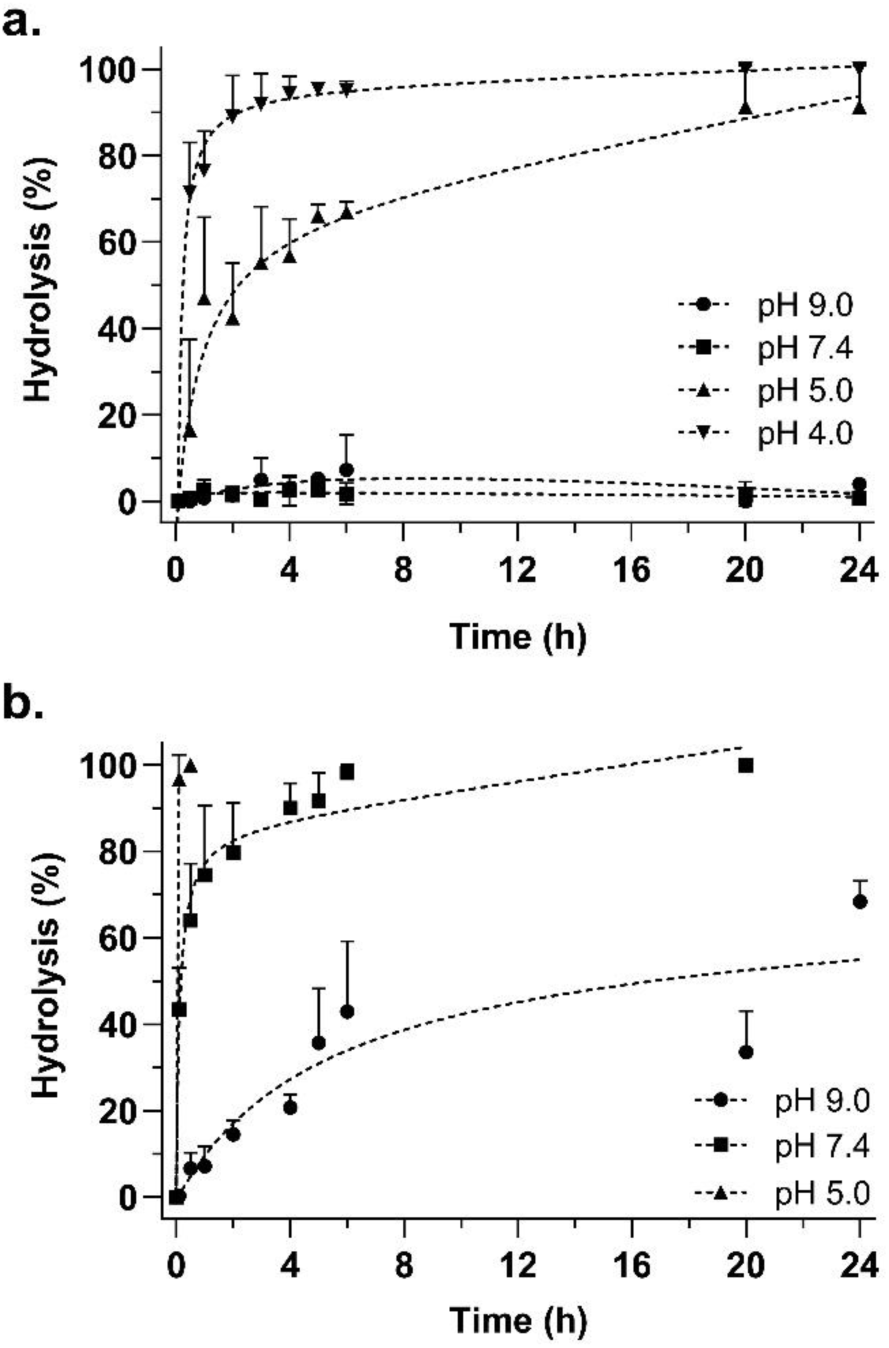
Hydrolysis of PEG-ma-Ale (a) and PEG-mma-Ale (b) cleavable conjugates at different pHs. PEG-ma-Ale showed excellent stability after 24 h incubation at both pH 9.0 and 7.4 (< 10% hydrolysis). A sustained Ale release was observed at pH 5.0, with ∼50% hydrolysis reached after about 3 h. At pH 4.0, the conjugate was almost completely cleaved (> 80%) within 1 h. On the other hand, PEG-mma-Ale showed rapid cleavage at pH 7.4 and 5.0 and slow amide cleavage at pH 9.0. Data are presented as mean + SD (n = 3).

### 3.2 CaP NPs preparation and characterization

PEG-ma-Ale stabilized CaP NPs were prepared by co-precipitation of calcium and phosphate ions in aqueous buffer, followed by quick addition of the Ale conjugate. Blank particles (not loaded with pDNA) had a hydrodynamic diameter of about 140 nm and a PDI of 0.1-0.2, while pEGFP encapsulation increased the particle size to around 170 nm (Table 1). In both cases, ζ-potential values were slightly negative and close to neutrality (−5 to −2 mV), which is in agreement with previous data for similar NPs [20]. The size difference between 10 and 20 µM PEG-ma-Ale was not significant both for blank and pDNA loaded CaP NPs. To confirm DNA entrapment, the gene loading efficiency was measured, and found to be of 95.4 ± 0.9% and 93.7 ± 1.0% at a PEG-ma-Ale concentration of 10 and 20 µM, respectively. Morphology of CaP NPs was investigated by TEM. CaP NPs displayed a spherical shape with a diameter of ca. 50 nm, forming clusters of several single units, with a smaller size compared to the hydrodynamic diameter measured *via* DLS (Fig. 3a). This finding correlates with previously reported data of similar systems [21], indicating that the CaP NPs may be present in suspension as small aggregates of single particles. No diffraction spots typical of crystalline materials were observed in the selected-area electron diffraction (SAED) (Fig. 3b), revealing that the particles were mainly composed of amorphous calcium phosphate [21]. Size, morphology and crystallinity degree of CaP nanoprecipitates are affected by the reaction conditions, such as initial Ca/P ratio, pH, presence of additives (e.g. ions, charged molecules and polymers), temperature and pressure [38,39]. When CaP NPs are produced by precipitation under super saturated conditions, amorphous CaP is generally formed first during the nucleation process, being the most kinetically favored CaP form [21,40,41]. For the *in vivo* studies (*vide infra*), the NPs suspension had to be concentrated to allow a sufficient dose of pDNA to be delivered. The particles could not be concentrated by centrifugation or ultrafiltration as aggregation and partial dissolution were observed. Therefore, the NPs were freeze-dried in the presence of a trehalose as a lyoprotectant. HEPES and TRIS buffers at lower ionic strength were used, to compensate for the increase in osmolality that occurred after concentration. pEGFP loaded CaP NPs prepared under “low osmolality” conditions had a hydrodynamic diameter of about 150 nm with a PDI of 0.15 (Fig. **S16**) and an osmolality of 139 mOsm/kg. Resuspension of the particles after lyophilisation resulted in a small but significant (p = 0.0480) increase in size to ∼200 nm. CaP NPs prepared with 10 µM PEG-ma-Ale were stable at RT for 3 days, whereas 15 and 20 µM PEG-ma-Ale increased particle stability to 7 and 15 days, respectively (Fig **S17**). While a higher concentration of PEGylated chelator was shown to be more efficient in preventing particle size growth [20], slow PEG-ma-Ale hydrolysis on the particle surface could explain the physical instability of the system upon long term storage. CaP NP stability in the presence of serum components was assessed by DLS at 37 °C. The average hydrodynamic diameter of PEG-ma-Ale stabilized CaP NPs did not significantly vary after 6 h incubation in DMEM containing 10 or 50% FBS, indicating good colloidal stability in biological media (Fig. 4a), at least under these non-diluted conditions. The addition of 10% FBS, resulted in a size increase from ∼160 to ∼220-240 nm, which was more pronounced at a lower PEG-ma-Ale feeding concentration (10 µM). This effect may be related to unspecific protein binding on the particle surface and/or more pronounced particle aggregation [21]. To assess the pH-dependent dissolution of the particles, PEG-ma-Ale stabilized CaP NPs were incubated at RT in 10 mM TRIS, and pH was slowly decreased by addition of 1N HCl. No significant change in size was initially observed in the pH range 6.8-7.4 (Fig. 4b). Upon acidification of the suspension buffer from pH 7.4 to pH 6.5, mimicking the early endosome environment, size of the particles dropped to ∼5 nm, indicating nearly complete dissolution.

**Table 1.** Physicochemical properties of PEG-ma-Ale coated CaP NPs. Means ± SD (n = 3-6).

| PEG-ma-Ale (μM) | Blank CaP NPs | | pEFGP loaded CaP NPs | |
| --- | --- | --- | --- | --- |
|  | 10 | 20 | 10 | 20 |
| Diameter (nm) | 138 ± 7 | 142 ± 4 | 170 ± 8 | 162 ± 4 |
| PDI | 0.19 | 0.14 | 0.105 | 0.115 |
| ζ-potential | -5 ± 1 | -1 ± 3 | -3 ± 2 | -3 ± 2 |

**Figure 3.**
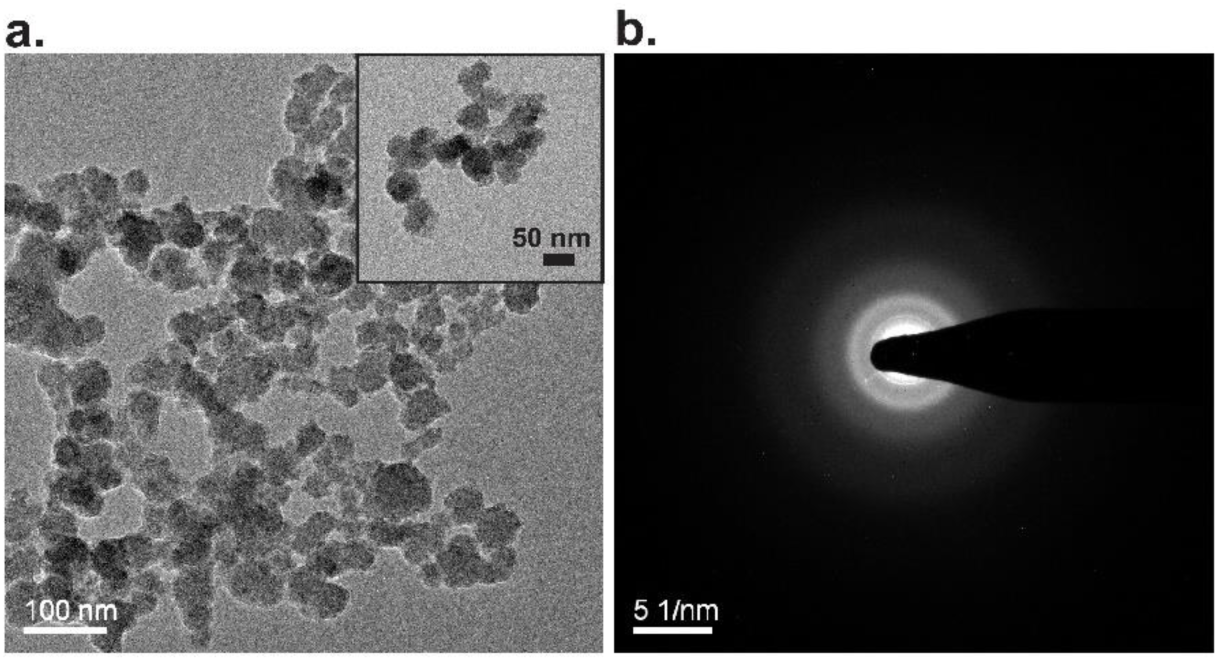
CaP precipitates formed spherical amorphous NPs. TEM micrograph (a) and SAED acquisition (b) of CaP NPs prepared with 10 µM PEG-ma-Ale.

**Figure 4.**
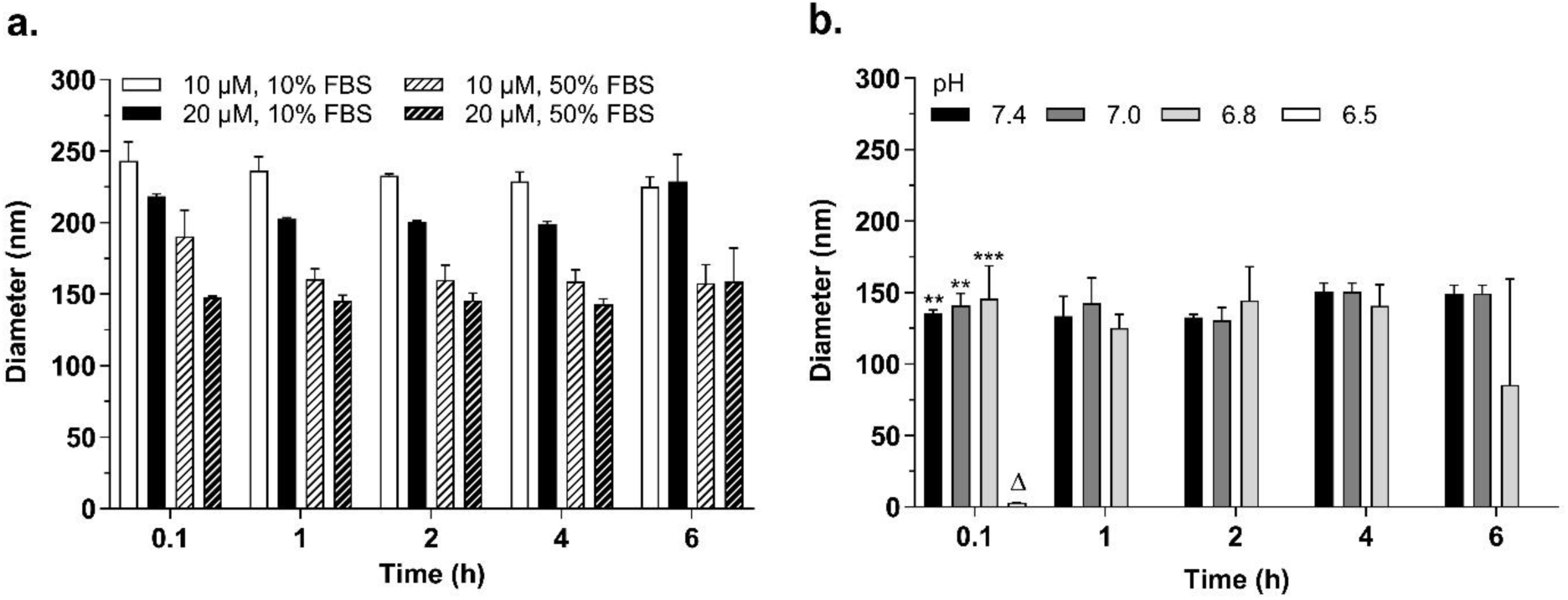
PEG-ma-Ale-chelated CaP NPs were stable in FBS-containing media, and quickly dissolved at mild acidic pH. The hydrodynamic diameter of PEGylated CaP NPs (PEG-ma-Ale 10 µM or 20 µM) was measured by DLS in the presence of 10 or 50% FBS (a). Size of CaP NPs was constant between pH 7.8 and 6.8, and the particles almost completely dissolved (Δ) upon acidification to pH 6.5 (b). Data are expressed as mean + SD (n=3). ** p ≤ 0.01 and *** p ≤ 0.001 *vs.* pH 6.5, 0.1 h.

### 3.3 Uptake of CaP NPs in 4T1 and J774.2 cells

To investigate the uptake of PEG-ma-Ale coated CaP NPs in 4T1 cancer cells and J774.2 macrophages, the particles were loaded with a fluorescein-labeled pDNA and incubated with the cells for 5 or 24 h in complete medium containing 10% FBS. X-tremeGENE™, a commercially available lipid-based transfection reagent, was used as positive control. Although the extent of cell-associated fluorescence was slightly different between the two cell lines, in both cases a modest uptake (∼20%) was observed after 5 h incubation compared to the positive control (∼70 %). A small but significant (p = 0.0043) lower uptake efficiency difference was found for the 20 µM PEG-ma-Ale formulation compared to the 10 µM one in 4T1 cells (Fig. 5a), which may be related to the higher surface PEG density. An increase of the incubation time to 24 h resulted in a 3-to 6-fold increase in particle internalization in 4T1 and J774.2 cells, respectively (Fig. 5b and 5c). In order to investigate whether particles sedimentation could affect CaP NPs cellular internalization, the same experiments were performed for the 24 h incubation time point with cells in the inverted configuration [42]. As expected, uptake efficiency of CaP NPs was significantly reduced in both cell lines in the inverted set up (cells on top), indicating that the internalization is dependent on the extent of particle sedimentation (Fig. 5b and 5c). We previously reported that the cellular uptake of CaP NPs in different cell lines was strongly influenced by their size, shape and surface PEG density [16,21]. Amorphous spherical particles of 250-280 nm were internalized by HepG2 cells at an almost 4-fold higher extent compared to similar NPs of 150 nm after 4 h incubation, possibly in part due to a larger sedimentation effect [21]. In addition, a higher surface PEG density generally reduces particles adhesion to cell and hinders their uptake [43]. For example, we reported that the uptake of 80 nm-CaP NPs stabilized with 30 or 40 µM PEGylated chelator was drastically decreased to around 10% compared to larger particles stabilized with a lower PEG concentration [21].

**Figure 5.**
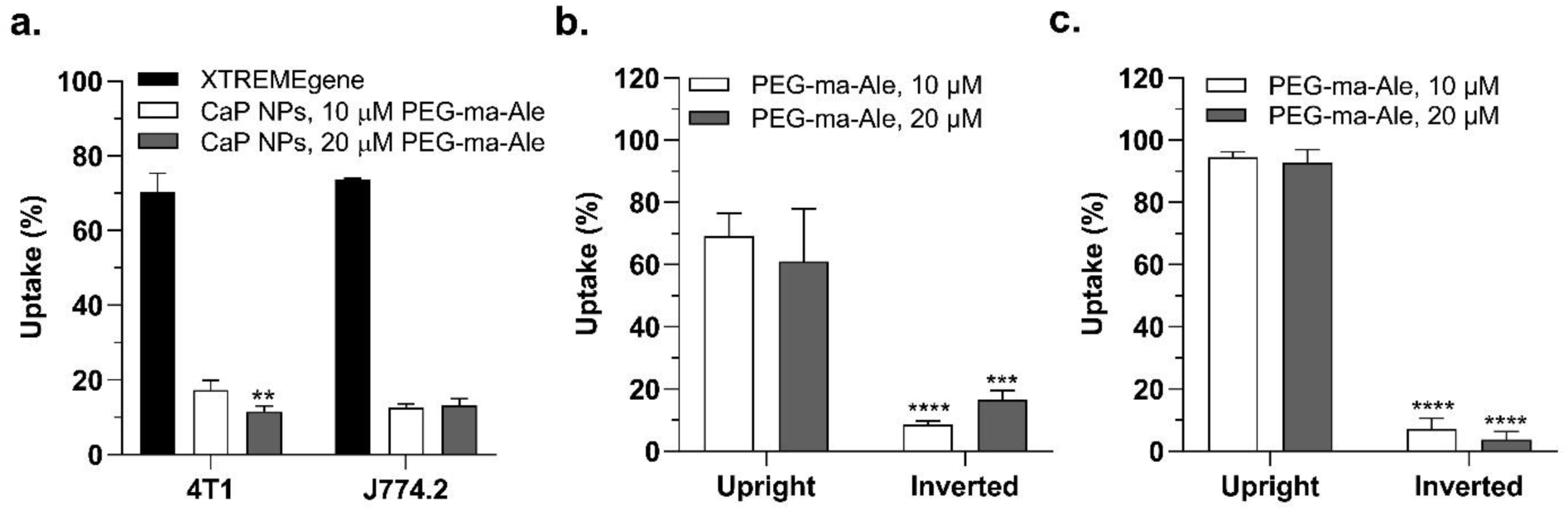
Cellular uptake of PEG-ma-Ale stabilized CaP NPs encapsulating a fluorescently labelled pDNA, after 5 h incubation with 4T1 and J774.2 cells (a), and after 24 h incubation in upright or inverted set-up with 4T1 (b) and J774.2 (c) cells. A 0.5-μg amount of DNA per well was applied and X-tremGENE™ was used as positive control. Data are expressed as mean + SD, n=3-6. ** p ≤ 0.01 *vs.* CaP NPs, 10 µM PEG-ma-Ale on 4T1 cells; *** p ≤ 0.001 and **** p ≤ 0.0001 *vs.* CaP NPs in the upright configuration at the same PEG-ma-Ale concentration.

### 3.4 Transfection efficiency of CaP NPs in 4T1 and J774.2 cells

The transfection ability of PEG-ma-Ale stabilized CaP NPs loaded with pEGFP plasmid was investigated by flow cytometry in 4T1 and J774.2 cell. CaP NPs were diluted in Opti-MEM supplemented with 10% FBS and exposed to cells for different incubation times (5, 24 and 48 h). X-tremeGENE™ was used as positive control. The EGFP expression in 4T1 cells increased with incubation time, ranging from a negligible (< 10%) transfection efficiency after 5 h cell exposure to CaP NPs, to about 50% EGFP positive cells after 48 h incubation, compared to ∼80% cells transfected with xTREMEgene™ (Fig. 6a and **S18**). This finding may correlate with a higher amount of CaP NPs internalized by cells for longer incubation times due to a sedimentation effect, as previously discussed. A lower transfection efficiency was observed for the particles coated with 20 µM of PEG-ma-Ale compared to 10 µM, which may be associated to the lower cellular uptake [21]. On the other hand, low or no transfection was observed for J774.2 macrophages incubated with PEG-ma-Ale stabilized CaP NPs (Fig. 6b). In both cell lines, few EGFP expressing cell were detected after 48 h incubation with the control uncoated CaP precipitates (Fig. **S19**). Discrepancies in the transfection efficiency of CaP NPs in different cell types were previously highlighted by Neuhaus et al. [44], investigating triple-shell CaP NPs (CaP–DNA–CaP– polyethylenimine). Despite the fact that multi-shell CaP NPs were rapidly taken up by all the cell lines used in their study, a lower transfection efficiency was achieved in 4T1 cells compared to HeLa cells, and no EGFP expressing cells were observed among monocytic cell lines. After CaP NPs internalization, endosomal escape of the cargo and DNA processing follow various intracellular mechanisms dependent on the cell type, which may eventually result in a different amount of protein expressed. A high transfection efficiency generally occurs in cells with a high proliferation rate, e.g., Hela, compared to cells that divide slower, such as 4T1 cells [45]. Moreover, due to their scavenging role monocytes and macrophages possess a high enzymatic activity which may result in DNA degradation before the genetic material reaches the nucleus [46]. We previously reported that CaP NPs stabilized with similar PEGylated chelators had a high transfection efficiency in HeLa cells (∼60% cells expressing GFP) [16]. The ability of our CaP NPs to transfect 4T1 cells was slightly lower compared to HeLa. On the contrary, their transfection efficiency in macrophages remained low.

**Figure 6.**
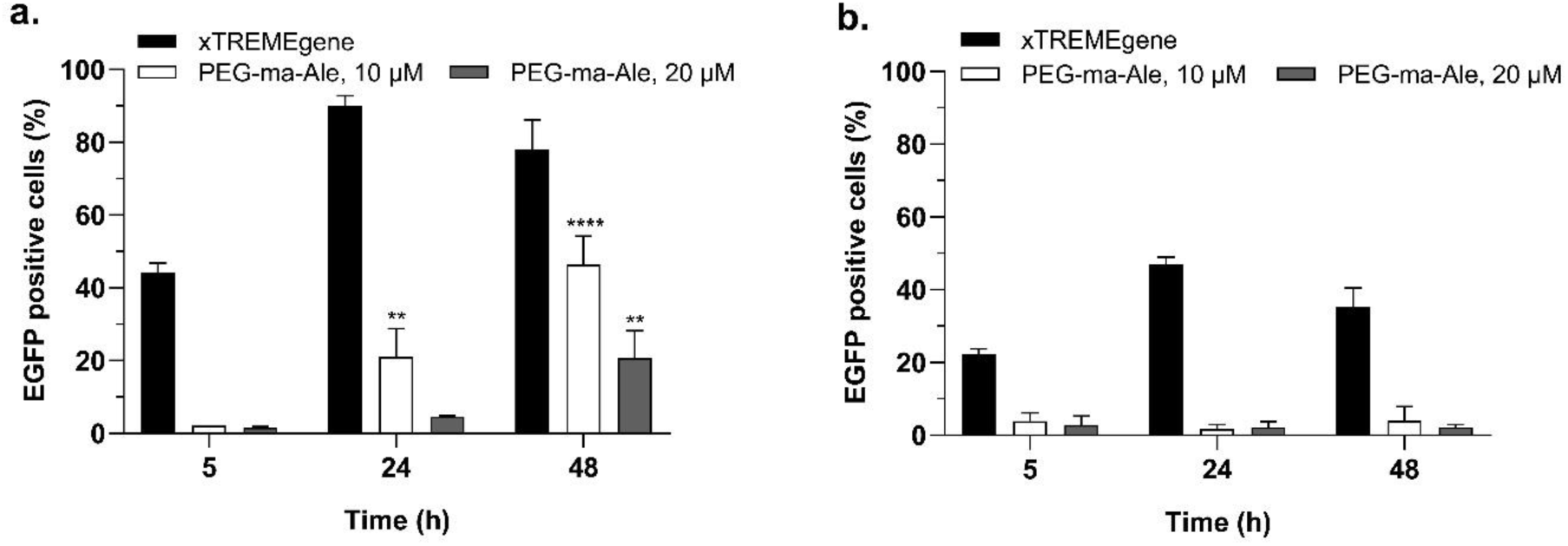
Influence of incubation time and PEG-ma-Ale concentration on transfection efficiency of CaP NPs in 4T1 (a) and J774.2 (b) cells in Opti-MEM containing 10% FBS. A prolongation of the incubation time from 5 to 48 h significantly increased the percentage of EGFP positive cells among 4T1 cells. In contrast, low CaP NPs transfection efficiency was observed in J774.2 cells at all tested conditions. Results are expressed as mean + SD (n=3-6). ** p ≤ 0.01 and *** p ≤ 0.0001 *vs.* 5 h incubation with CaP NPs at the same PEG-ma-Ale feeding concentration.

### 3.5 Cytotoxicity

The cytotoxicity of PEG-ma-Ale stabilized CaP NPs was evaluated on 4T1 and J774.2 cells, following exposure to the particles for increased PEG-ma-Ale concentrations (0.5, 1, 2, 4 and 5 µM, based on the corresponding Ale molar concentrations) and incubation times (5, 24 and 48 h). As a control, CaP NPs coated with the non-cleavable PEG-a-Ale derivative were used. These particles had physicochemical properties comparable to those of PEG-ma-Ale stabilized CaP NPs (Table **S1**), and their characteristics were previously reported [20]. On J774.2 macrophages, PEG-ma-Ale coated CaP NPs induced a time- and concentration-dependent cytotoxic effect, displaying a significant decrease of cell viability (∼60-70%) after 24 h exposure to concentrations of conjugated Ale ≥ 0.5 µM (Fig. 7a). After 48 h incubation and PEG-ma-Ale concentrations ≥ 1 µM, J774.2 viability dropped to less than 20%. Incubation with free PEG-ma-Ale resulted in a moderate but significant decrease of cell viability (∼20-35%) at high conjugate concentrations (≥ 4 µM) (Fig. **S20**). Conversely, CaP NPs stabilized with the non-cleavable PEG-a-Ale derivative did not display cytotoxicity on macrophages under similar conditions (Fig. 7b). Uncoated CaP precipitates (no Ale) had no effect on macrophages viability at the same dilution factor tested for the other CaP NPs formulations (Fig. **S21**). As shown in Fig. 7c, PEG-ma-Ale associated with CaP NPs was substantially less cytotoxic towards 4T1 cells compared to macrophages. Treatment with nitrogen-containing BPs was reported to affect *in vitro* viability of different cell lines, primarily osteoclasts, monocytes and macrophages. For free Ale, a dose-dependent cytotoxic effect in monocytic and cancer cell lines was observed for rather high concentrations (20-500 µM, depending on the cell line) [47–50]. However, adsorption or encapsulation of the BP in nanoparticulate systems were shown to drastically decrease the half maximal inhibitory concentration of Ale, as they facilitated the cellular uptake of the drug [50–53]. BPs induction of apoptosis is correlated to inhibition of the mevalonate pathway, which leads to reduced biosynthesis of isoprenoid compounds that are essential for protein prenylation [49]. Prenylated GTPases, such as Rap, Rho, Rac, Rab and Ras, are involved in the regulation of cytoskeletal organization, cell morphology, endocytosis and apoptosis. Therefore, insufficient GTPases prenylation results in impaired cell function and cell death. To further examine whether cytotoxicity of CaP NPs was mediated by inhibition of FPPS activity, we measured the level of an unprenylated GTPase, Rap1A, after 24 h incubation with CaP NPs coated with PEG-ma-Ale or the more stable control PEG-a-Ale [20]. Accumulation of unprenylated Rap1A was observed in both cell lines treated with CaP NPs bearing the pH-sensitive derivative, at concentrations of conjugated Ale ≥ 0.5 µM (Fig. 7d and 7e). The effect was stronger on macrophages, for which the particles uptake was slightly higher compared to 4T1 cells (Fig. 5b and 5c). On the other hand, the unprenylated protein was not detected in lysates from cells exposed to PEG-a-Ale chelated CaP NPs. These findings indicate that the cytotoxicity associated to PEG-ma-Ale stabilized CaP NPs in both cell lines was correlated to the inhibition of protein prenylation, which is the major mechanism induced by Ale that ultimately triggers cell apoptosis. Since the protonated form of the Ale nitrogen is required for the binding to FPPS [54], we can hypothesized that pH-driven hydrolysis of PEG-ma-Ale allows for a greater intracellular release of Ale. In comparison, no detectable accumulation of unprenylated Rap1A after exposure to PEG-a-Ale stabilized CaP NPs may indicate insufficient cleavage of Ale from the stable conjugate [20].

**Figure 7.**
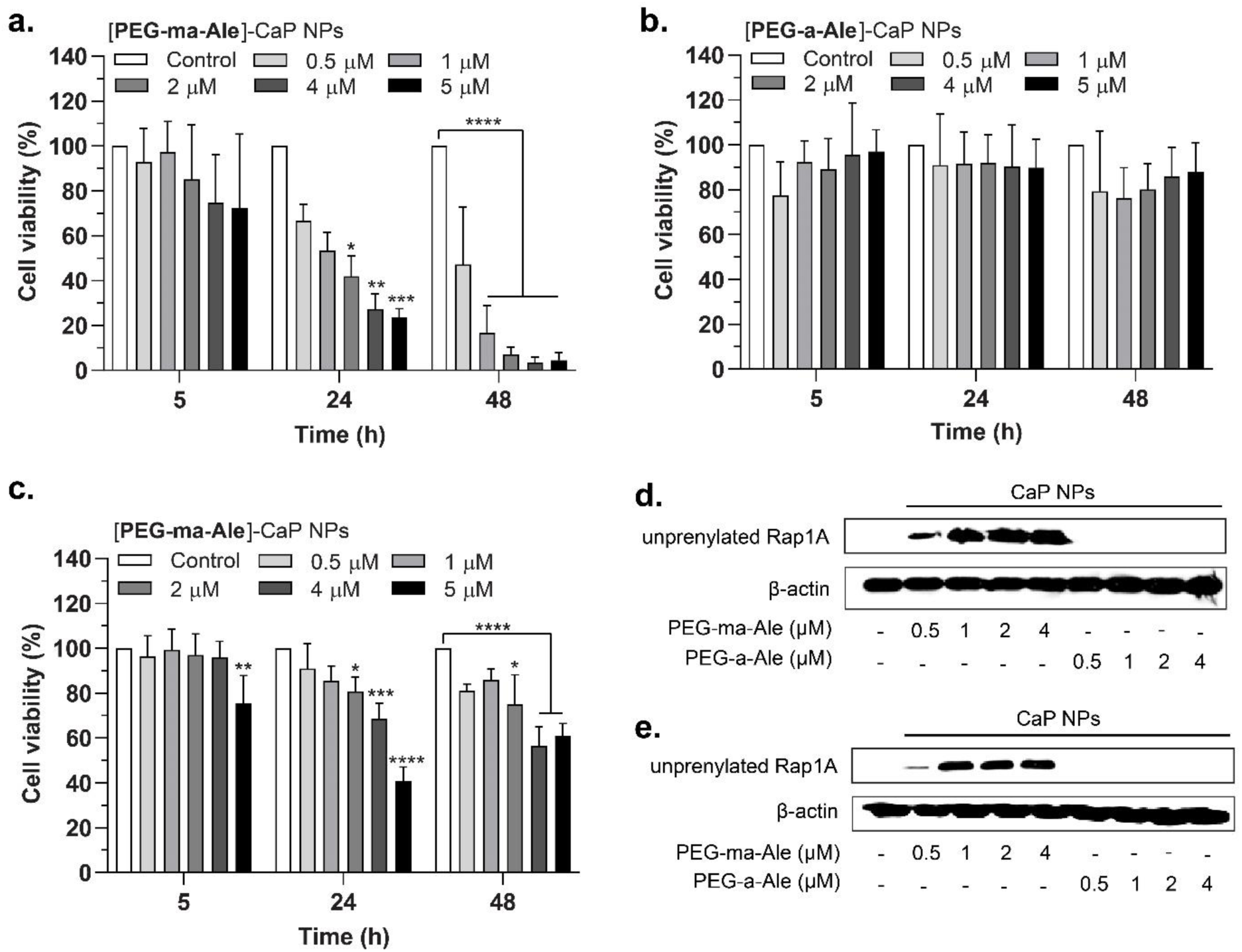
PEG-ma-Ale stabilized CaP NPs affected cell viability of macrophages and cancer cells and induced accumulation of unprenylated Rap1A, via inhibition of the mevalonate pathway. Cell viability after 5, 24 or 48 h incubation of PEG-ma-Ale stabilized CaP NPs (a) and PEG-a-Ale stabilized CaP NPs (b) with J774.2 cells. In the same way, 4T1 cell viability was evaluated after exposure to PEG-ma-Ale coated CaP NPs (c). Data are expressed as mean + SD of three independent experiments performed in triplicate. * p ≤ 0.05; ** p ≤ 0.01; *** p ≤ 0.001; **** p ≤ 0.001 *vs*. control wells (not treated). Western blot analyses were performed to examine unprenylated Rap1A accumulation in J774.2 (d) and 4T1 (e) cell lysates after 24 h incubation with CaP NPs coated with PEG-ma-Ale or PEG-a-Ale. Representative Western blot of 3 independent experiments.

**Figure 8.**
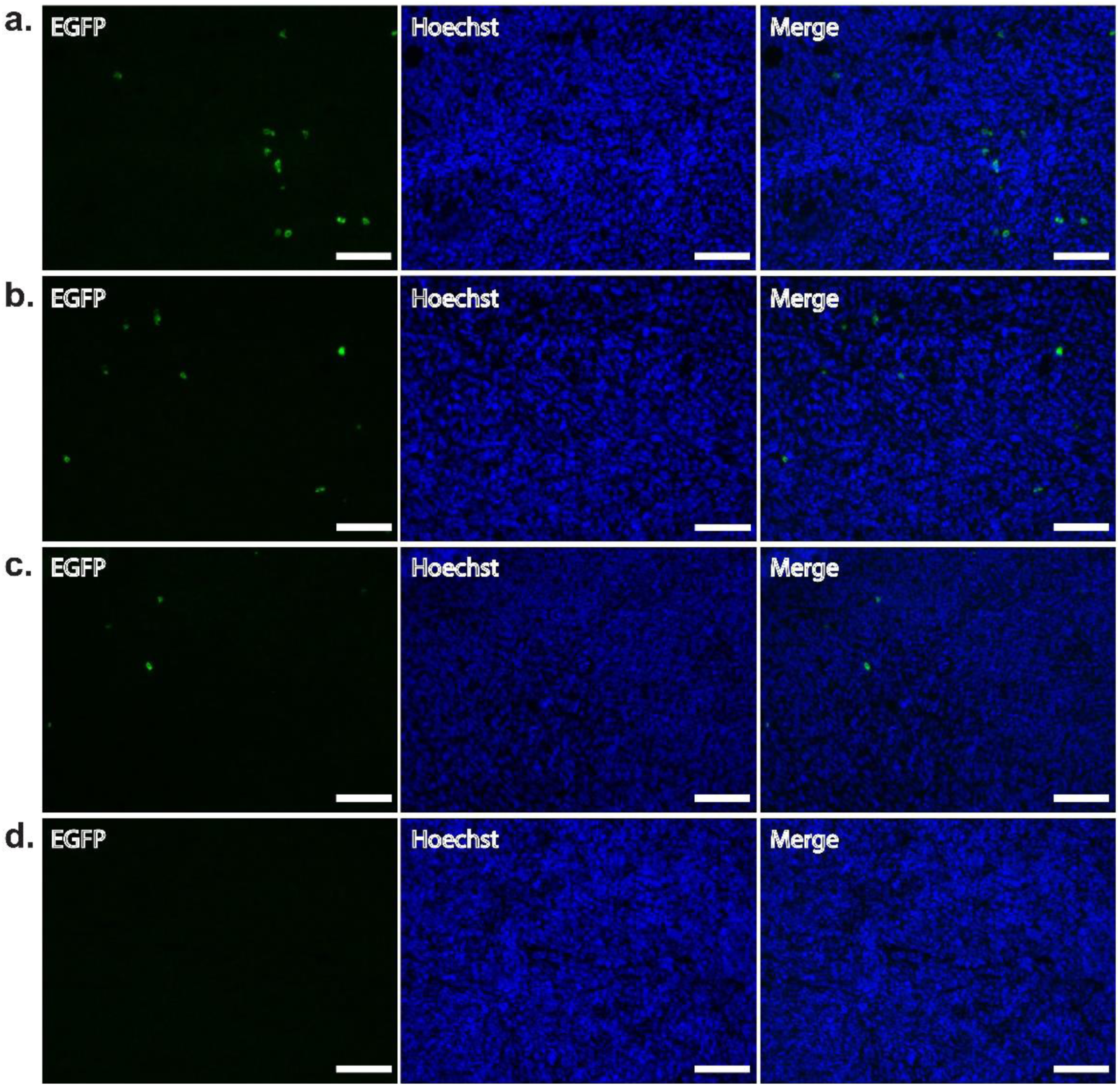
Representative fluorescence micrographs of histological analysis of tumour frozen sections after administration of PEG-ma-Ale stabilized CaP NPs (a) *in vivo*-jetPEI^®^ (b) and naked pEGFP (c). A solution of 5% (*m/v*) glucose was used as negative control (d). The formulations were injected intratumorally in 4T1 xenografts in BALC/c mice every other day for one week. Samples were isolated 24 h after the last injection. For EGFP visualization, the tissue slices were stained with ATTO488-GFP Booster. Scale bar represents 100 µm.

### 3.6 *In vivo* transfection efficiency

Since PEG-ma-Ale stabilized CaP NPs displayed good *in vitro* transfection efficiency in 4T1 cancer cells, we investigated their ability to deliver *in vivo* pEGFP to xenografted 4T1 breast cancer tumours, following intratumoral injection, to maximize the amount of pDNA delivered to the tumour. Due to hyperosmolality of the reconstituted CaP NPs after concentration, a small volume of CaP NPs formulation (20 µL) was slowly injected in multiple spots of the tumor, to avoid pain and tissue damage [28,55]. The transfection efficiency study was performed by administration of three single doses of pEGFP loaded CaP NPs (each dose consisting of 2 µg), every other day over one week. *In vivo*-jetPEI^®^, a commercial cationic polymer-based agent for *in vivo* transfection, was used as a positive control. A low GFP expression was found in the tumors injected with PEG-ma-Ale stabilized CaP NPs, which was similar to that of *in vivo*-jetPEI^®^. On the other hand, a lower transfection was observed after administration of the naked plasmid, suggesting that pDNA incorporation in NPs increases the degree of DNA transfer efficiency across the cell membrane. However, the low *in vivo* transfection efficacy of the PEG-ma-Ale coated CaP NPs could be potentially explained by a lower pDNA dose compared to those found in the literature [56], a lower particle uptake *in vivo* (due to the absence of a sedimentation effect) and/or to the premature CaP dissolution (and subsequent extracellular pDNA release) that may occur under dilute conditions [57]. Moreover, particles aggregation at the injection site and/or poor diffusion through the extracellular matrix could also have impaired the transfection ability of the system.

## Conclusions

In the present study, an acid-cleavable PEG-Ale conjugate was used to generate stable CaP NPs loaded with pEGFP and allow the combined cellular uptake of the two poorly permeable compounds. *In vitro*, the pH responsive nanocarrier exhibited good transfection efficiency in cancer cells and a strong cytotoxic effect on macrophages with potent inhibition of the mevalonate pathway, probably due to efficient endo/lysosomal cleavage of the BP from the particle coating. By providing an efficient approach for the combined delivery of drugs, further anticancer applications may be envisaged, *i.e.* replacing the model pDNA with therapeutic nucleic acids, if the *in vivo* transfection efficiency is improved. The introduction of a selected targeting ligand on the particles surface could boost their uptake in cancer cells or tumour associated macrophages, overcoming the negative effect of the PEG layer on internalization. While particle dissolution under mildly acidic condition allows for the complete discharge of the payload, further optimization strategies are needed to improve the stability of CaP NPs, which tend to dissolve under highly diluted conditions, such as those encountered *in vivo*. For example, particle dissolution may be tuned varying the crystalline phase of the nanomaterial [58], investigating hydrothermal treatments or synthetic routes that allow for a precise control over the Ca/P ratio [59].

## Acknowledgments

The authors wish to thank Catherine Cailleau at Institut Galien Paris-Sud, UMR8612, Université Paris-Sud, for the technical assistance in animal experiments. The Scientific Center for Optical and Electron Microscopy (ScopeM) of ETH Zurich is gratefully acknowledged for use of the facility, and especially Dr. Fabian Gramm for TEM and SAED analyses and Dr. Gabriella Éva Bodizs and Joachim Hehl for assistance with the cryostat and with EGFP staining. Antonia Schantl is acknowledged for proofreading of the manuscript. This work was financially supported by the State Secretariat for Education, Research and Innovation (SERI) in the context of the European Union’s HORIZON 2020 Programme (Marie Curie ITN European NABBA project, no. 642028) and a Natural Sciences and Engineering Research Council of Canada Discovery Grant (RGPIN-2015-05364). The authors declare no conflict of interest.

