## Supporting information for "Dual delivery of nucleic acids and PEGylated-bisphosphonates *via* calcium phosphate nanoparticles"

### Instrumentation

Proton nuclear magnetic resonance ($^1$H-NMR) spectra were acquired on a Bruker Av400 spectrometer (Bruker BioSpin, Fällanden, Switzerland) operating at 400 MHz.

### S1. Synthesis of PEG-Ale chelators for CaP NPs

#### *S1.1 Synthesis of PEG-OTs*

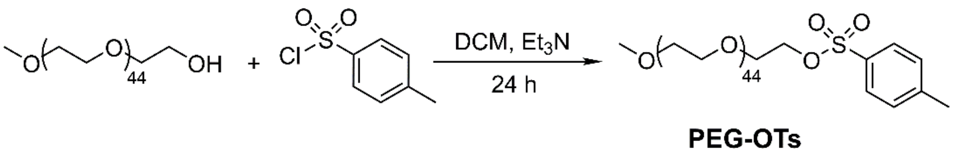

The synthesis of PEG-azide (PEG-$N_3$) for copper-catalyzed azide-alkyne cycloaddition was adapted from a previous protocol [1]. Briefly, mPEG-OH ($M_n \approx 2000$ Da) (5 g, 2.50 mmol, 1 eq.) was dissolved in 15 mL anhydrous dichloromethane (DCM). A fresh solution of toluenesulfonyl chloride (1.906 g, 10.00 mmol, 4 eq.) and triethylamine ($Et_3N$) (1.57 mL, 11.25 mmol, 4.5 eq.) in 15 mL DCM was added dropwise to the PEG solution under vigorous stirring, at 0 °C and under nitrogen atmosphere. After 24 h, the reaction mixture was concentrated by rotary evaporation, dissolved in warm (37 °C) ethanol, then cooled down in ice, and the precipitate was collected by centrifugation (3x). Subsequently, the crude product was dissolved in DCM and isolated by precipitation (3x) in diethyl ether ($Et_2O$). The precipitate was collected by centrifugation and dried under vacuum to obtain the final product (PEG-OTs) (3.83 g, 70.7 %). **$^1$H-NMR** (400 MHz; $CDCl_3$): δ 7.74 (m, 2H), 7.29 (m, 2H), 3.77-3.39 (m, 200H), 3.31 (s, 3H), 2.39 (s, 3H) (Fig. **S1**).

#### *S1.2 Synthesis of PEG-$N_3$*

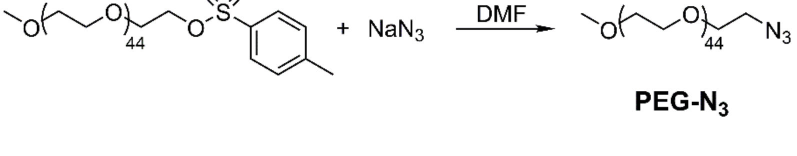

PEG-OTs (2 g, 0.92 mmol, 1 eq.) and sodium azide ($NaN_3$) (237 mg, 3.65 mmol, 4 eq.) were suspended in 8 mL anhydrous *N,N*-dimethylformamide (DMF) and stirred overnight under nitrogen atmosphere, at

RT. Then, the reaction mixture was poured into a brine solution and extracted with DCM (3x). The organic layers were combined and washed once with ultrapure water, dried over sodium sulphate ($Na_2SO_4$) and filtered. The solvent was evaporated and the crude product was isolated by precipitation in $Et_2O$ (3x). The precipitate was collected by centrifugation and dried under vacuum to obtain the final product (PEG-$N_3$) (1.44 g, 77 %). **$^1$H-NMR** (400 MHz; $CDCl_3$): δ 3.82-3.45 (m, 200H), 3.38-3.36 (m, 5H) (Fig. **S2**).

#### *S1.3 Synthesis of Compound 1*

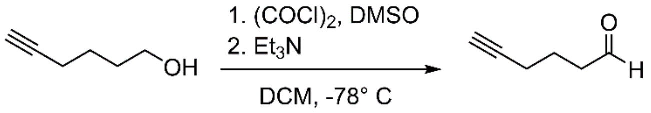

**1**

Compound **1** (5-hexan-1-al) was obtained following a reported procedure [2] with minor modifications. Oxalyl chloride (($(COCl)_2$, 1.29 mL, 15.28 mmol, 1 eq.) was dissolved in 37.5 mL DCM and the solution was cooled down to -78°C. Dimethyl sulfoxide (DMSO, 2.38 mL, 33.61 mmol, 2.2 eq.) was mixed with 8 mL DCM, then added to $(COCl)_2$ over a period of 20 min, and stirred for 30 min. 5-hexan-1-ol (1.68 mL, 15.28 mmol, 1 eq.) was dissolved in 20 mL DCM and the solution was added to the reaction mixture dropwise, over 10 min. The reaction was carried out for 1 h at -78°C. Then, $Et_3N$ (5.32 mL, 38.18 mmol, 2.5 eq.) was added and the reaction mixture was stirred for 1 h at -78°C and then allowed to warm to 10 °C over an additional hour. Forty mL of ultrapure water were added to the reaction mixture for the extraction. The water phase was separated and acidified with hydrochloric acid (HCl), and back extracted with DCM (3 x 20 mL). The organic extracts were washed with 1% HCl in water saturated with NaCl (6 x 20 mL) followed by addition of 5 % sodium bicarbonate solution (2 x 11 mL). The organic extracts were washed with brine (2 x 5 mL), dried over $Na_2SO_4$ and filtered. The organic solvent was removed by rotary evaporation (temperature of the water bath below 30 °C), and compound **1** was obtained as a yellow oil (1.16 g, 79.5%). **$^1$H-NMR** ($CDCl_3$): δ 9.81 (q, J = 1.4 Hz, 1H); δ 2.60 (td, J = 7.2, 1.3 Hz, 2H); δ 2.27 (tdd, J = 6.9, 2.7, 1.2 Hz, 2H); δ 1.98 (td, J = 2.7, 0.8 Hz, 1H), δ 1.90-1.81 (m, 2H) (Fig. **S3**).

### *S1.4 Synthesis of Compound 2*

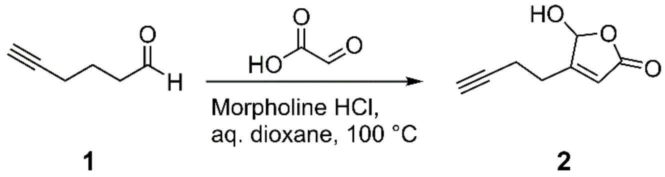

Compound **2** was obtained following a reported procedure [3], with minor modifications. Morpholine (7.23 mL, 83.83 mmol) was dissolved in about 60 mL $Et_2O$, under stirring. HCl 37% (6.84 mL, 83.83 mmol) was added dropwise to morpholine. $Et_2O$ and water were removed by rotary evaporation and a white solid was obtained. The product was washed 4 times with 1,4-dioxane, followed by 4 times washing with $Et_2O$. $Et_2O$ was removed by rotary evaporation, and the white solid was dried under vacuum, to yield the morpholinium·HCl salt (8.32 g, 80.3%). Glyoxylic acid monohydrate (3.729 g, 40.52 mmol, 1 eq.) was dissolved in 75 mL 1,4-dioxane, and morpholinium·HCl salt (5.167 g, 42.54 mmol, 1.05 eq.) was added to the reaction mixture. Ultrapure water was added dropwise until the complete dissolution of the morpholine salt. Compound **1** (4.09 mL, 40.52 mmol, 1 eq.) was dissolved in 12 mL 1,4-dioxane and added dropwise to the reaction mixture. After 3 h stirring at RT, the flask was equipped with a reflux condenser and the reaction mixture was heated up to 100 °C for 14 h. Then, the reaction mixture was cooled down and dioxane was removed by rotary evaporation. The isolated material was transferred in a separation funnel with 10 mL $Et_2O$ and 10 mL ultrapure water, the organic layer was collected, and the aqueous layer was extracted twice with $Et_2O$. The organic extracts were washed with brine, dried over $Na_2SO_4$, filtered and concentrated under vacuum. The crude product was purified by silica chromatography, using a 20 → 40% ethyl acetate/n-hexane elution gradient. Solvents were removed by rotary evaporation to isolate compound **2** (2.01 g, 32.6%). **$^1$H-NMR** ($CDCl_3$): δ 6.08 (d, $J$ = 0.8 Hz, 1H); δ 6.00 (td, $J$ = 1.6, 0.9 Hz, 1H); δ 2.79 – 2.67 (m, 1H), δ 2.66 – 2.56 (m, 1H); δ 2.56 – 2.49 (dddd, $J$ = 8.7, 7.5, 4.7, 1.9 Hz, 2H); δ 2.05 – 2.02 (t, $J$ = 2.6 Hz, 1H) (Fig. **S4**).

### *S1.5 Synthesis of Compound 3*

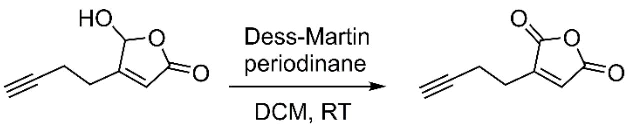

**2** **3**

Dess-Martin periodinane (313.8 mg, 0.74 mmol, 1.05 eq.) was suspended in 2.5 mL DCM. Compound **2** (107.5 mg, 0.71 mmol, 1 eq.) was dissolved in 1 mL DCM and added dropwise to the Dess-Martin periodinane suspension over a period of 10 min. The reaction mixture was stirred for 3 h at RT. Fifteen mL of a 1:1 (*v/v*) $NaHCO_3/Na_2SO_3$ saturated aqueous solution were added to the reaction mixture for quenching, and the suspension was stirred for 15 min. The aqueous phase was extracted with DCM (3x), the organic extracts were washed with brine, dried over $Na_2SO_4$, filtered and concentrated under vacuum. The crude product was purified by silica chromatography, using a 96:4 → 92:8 DCM/acetone elution gradient. Solvents were removed by rotary evaporation to isolate compound **3** (40.9 mg, 38.53 %). **$^1$H-NMR** ($CDCl_3$): δ 6.81 (s, 1H); δ 2.76 (m, 2H); δ 2.59 (m, 2H); δ 2.07 (t, 1H) (Fig. **S5**).

### *S1.6 Synthesis of Compound 4*

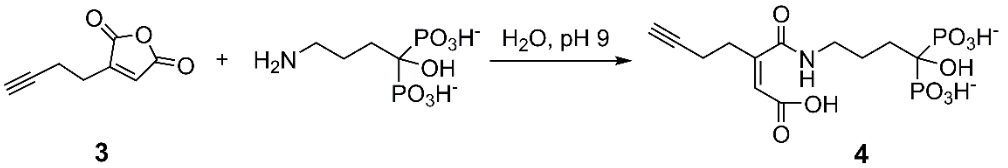

**3** **4**

Compound **3** (23.6 mg, 0.16 mmol, 1 eq.) was mixed with 0.2 mL ultrapure water. Alendronate (Ale) sodium trihydrate (76.21 mg, 0.23 mmol, 1.5 eq.) was dissolved in 0.4 mL 1N NaOH, and the solution was diluted to 0.7 mL with ultrapure water. Compound **3** was mixed with Ale solution and vortexed vigorously. The pH was adjusted to 9.0 by dropwise addition of 1N NaOH. The final pale yellow solution was stirred for 1 h on ice, then 1 h at RT. The crude compound **4** was not further purified and was characterized by $^1$H NMR spectroscopy. **$^1$H-NMR** ($D_2O$): δ 5.92 (s, 1H); δ 3.23 (m, 2H); δ 2.48 (m, 2H); δ 2.40 (m, 2H), δ 1.99-1.78 (m, 2H) (Fig. **S6**).

#### *S1.7 Synthesis of PEG-ma-Ale*

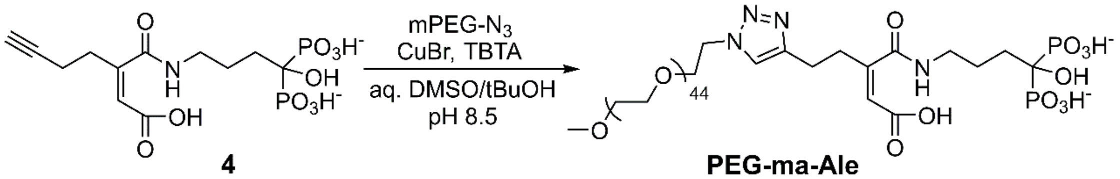

4 PEG-ma-Ale

PEG-ma-Ale was obtained by conjugation of compound **4** and PEG-$N_3$ *via* copper-catalysed azide-alkyne cycloaddition [4]. PEG-$N_3$ (63.58 mg, 31.18 µmol, 0.5 eq.) was dissolved in 0.6 µL 2 mM $Na_2HPO_4$, pH 8.5. Crude compound **4** (containing 24.8 mg, 62.75 µmol, 1 eq. of conjugated compound **3**) was dissolved in 1 mL of 2 mM $Na_2HPO_3$, pH 8.5 and added to the PEG-$N_3$ solution, under vigorous stirring. Click solution C (30 µL) was added to the reaction mixture to initiate the click reaction. This solution was freshly prepared by mixing click solution A and click solution B (1/2, *v/v*). Click solution A was freshly prepared by dissolving copper(I) bromide (CuBr) (0.8 mg, 5.58 µmol) in DMSO/*tert*-butanol (tBuOH) (70 µL, 3/1 *v/v*). Click solution B was prepared by dissolving tris[(1-benzyl-1H-1,2,3-triazol-4-yl)methyl]amine (TBTA) (2.7 mg, 5.08 µmol) in 100 µL DMSO/tBuOH (3/1 *v/v*). The reaction mixture was stirred for 3 h at RT. Then, insoluble salts were isolated by centrifugation (2 min, RT, 12,000 *x g*), and the final product was purified from free Ale by size exclusion chromatography, using a Sephadex G-25 PD-10 column and 10 mM $Na_2CO_3$, pH 9.0 as eluent. Fractions containing the PEG derivative were identified *via* cobalt thiocyanate staining [5], and the positive fractions were further purified by size exclusion chromatography as previously described (6x). The final product was lyophilized (PEG-ma-Ale, 64.7 mg 85.73%) and subsequently dissolved in 20 mM $Na_2HPO_4$ buffer, pH 7.4 in $D_2O$ and characterized by $^1$H-NMR spectroscopy. **$^1$H-NMR** ($D_2O$): δ 7.89 (s, 1H); δ 5.85 (s, 1H); δ 4.61 (m, 2H); δ 3.98 (m, 2H), δ 3.90-3.52 (m, 200H), δ 3.39 (m, 2H), δ 3.24 (m, 2H), δ 2.91 (m, 2H), δ 2.67 (m, 2H), δ 19.6 – 1.84 (m, 4H) (Fig. **S7**).

#### *S1.8 Synthesis of Compound 5*

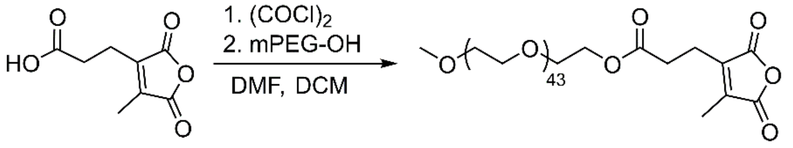

**5**

2,5-dihydro-4-metyl-2,5-dioxo-3-furanpropanoic acid (100 mg, 0.54 mmol, 1 eq.) was dissolved in 2 mL anhydrous DCM and placed in ice bath. $(COCl)_2$ (69.85 µL, 0.81 mmol, 1.5 eq.) was added to the reaction mixture, followed by dropwise addition of DMF (3.96 µg, 54.3 µmol, 0.1 eq.). The solution was stirred at 0 °C for 40 min, then at RT for 2 h. Toluene was added to the reaction mixture, which was then concentrated by rotary evaporation. The crude product was dissolved in 2 mL DCM. mPEG-OH ($M_n \approx 2000$ Da) (869 mg, 0.43 mmol, 0.8 eq.) was dissolved in 2 mL DCM, followed by addition of $Et_3N$ (83.25 µL, 60.44 mg, 0.60 mmol, 1.1 eq.). The reaction mixture containing PEG was added dropwise to the anhydride solution and stirred at RT for 1 h. Then, 10 mL of DCM and 10 mL of 1N HCl were added to the crude product, and the aqueous phase was extracted with DCM (3x). The organic fractions were collected, washed with brine, dried over $Na_2SO_4$ and filtered. The crude product was dissolved in DCM and precipitated in $Et_2O$ (3x). Compound **5** was isolated by filtration and dried under vacuum for 5 days (619.4 mg, 66.8%). **$^1$H-NMR** ($CDCl_3$): δ 4.2 (m, 2H); δ 3.79-3.44 (m, 200H); δ 3.35 (m, 3H); δ 2.72 (m, 4H), δ 2.10 (m, 3H) (Fig. **S8**).

#### *S1.9 Synthesis of PEG-mma-Ale*

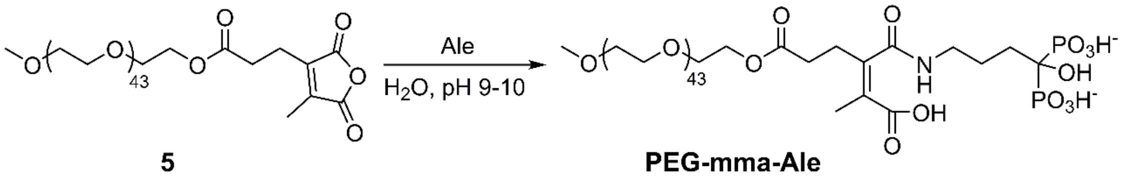

**5** **PEG-mma-Ale**

Ale sodium trihydrate (4.47 mg, 13.75 µmol, 1 eq.) was suspended in 150 µL 20 mM $Na_2CO_3$ buffer, pH 9.0, followed by addition of 10 µL 1N NaOH, to dissolve the salt. Compound **5** (30 mg, 13.75 µmol, 1 eq.) was dissolved in 200 µL of 20 mM $Na_2CO_3$, pH 9.0, and quickly added to the Ale solution, under vigorous stirring, in ice bath. pH was adjusted to 9-10 by dropwise addition of 1N NaOH. The reaction mixture was stirred for 30 min in ice bath, followed by 1 h at RT. The crude product was purified by size exclusion chromatography with a Sephadex G-25 PD-10 size exclusion column using 20 mM $Na_2CO_3$, pH 10 as eluent. PEG-containing fractions were identified *via* cobalt thiocyanate staining [5] and lyophilized to obtain the final product (PEG-mma-Ale, 16 mg, 48.3%). PEG-mma-Ale was dissolved in 20 mM $Na_2CO_3$, pH 9.5 in $D_2O$ and characterized by $^1$H-NMR. **$^1$H-NMR** ($D_2O$): δ 4.29 (m, 2H); δ 3.91 - 3.53 (m, 200H); δ 3.4 (m, 3H); δ 3.21 (m, 2H), δ 2.61-2.52 (m, 4H), δ 1.93 – 1.81 (m, 7H) (Fig. **S9**).

#### *S1.10. Synthesis of PEG-a-ALE*

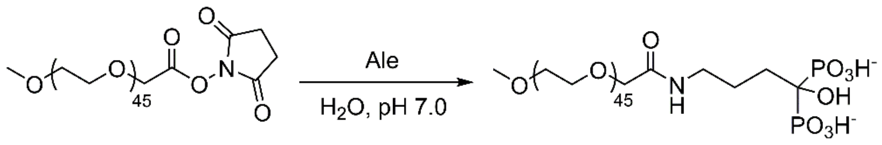

**PEG-a-Ale**

PEG-a-ALE was synthetized according to a previously reported protocol [5], with minor modifications. Ale sodium trihydrate (400 mg, 1.23 mmol) was dissolved in 1 mL 2N NaOH, followed by addition of 800 µL of ultrapure $H_2O$. pH was adjusted to 7 by dropwise addition of 1N HCl. Methoxy-PEG succinimidyl carboxymethyl ester ($M_n \approx 2000$ Da) (30 mg, 13.8 µmol) was dissolved in 200 µL ultrapure $H_2O$ and then added dropwise to the Ale solution, under vigorous stirring. The reaction mixture was stirred in ice for 1 h, followed by 1 h at RT. The compound was purified by size exclusion chromatography with a Sephadex G-25 PD-10 size exclusion column using ultrapure water as eluent. Fractions containing the PEG derivative were identified *via* cobalt thiocyanate staining [5]. Free PEG was removed by anion exchange using a column filled with Dowex 1×2, 200-400 mesh. As elution buffer, $Na_2HPO_3$ (10, 50, and 200 mM) was used. Fractions containing only conjugated PEG were collected, desalted using a Sephadex G-25 PD-10 size exclusion column (eluent ultrapure water) and finally freeze-dried to obtain PEG-a-Ale (15.3 mg, 48.2%). **$^1$H-NMR** ($D_2O$): δ 9.81 (q, J = 1.4 Hz, 1H); δ 2.60 (td, J = 7.2, 1.3 Hz, 2H); δ 2.27 (tdd, J = 6.9, 2.7, 1.2 Hz, 2H); δ 1.98 (td, J = 2.7, 0.8 Hz, 1H), δ 1.90-1.81 (m, 2H) (Fig. **S10**).

### S.2 Supporting results

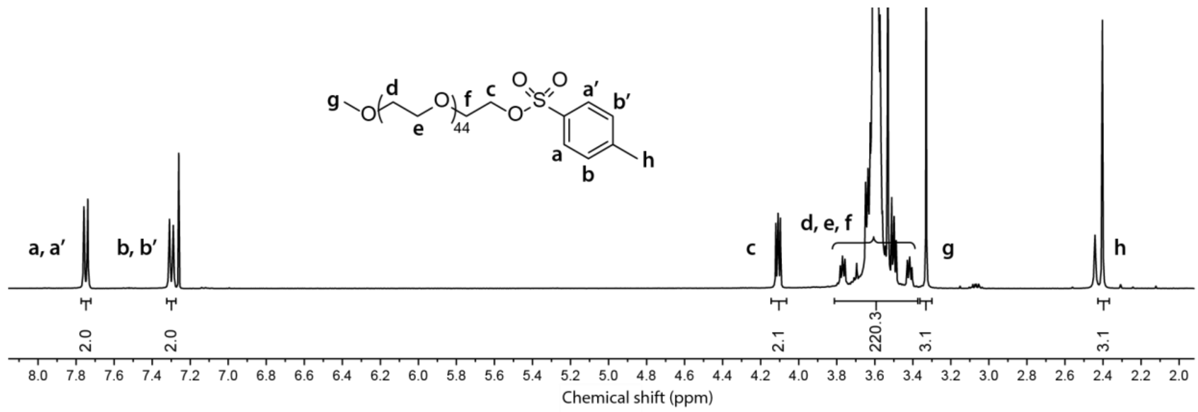

**Figure S1.** $^{1}$H-NMR spectrum of PEG-OTs.

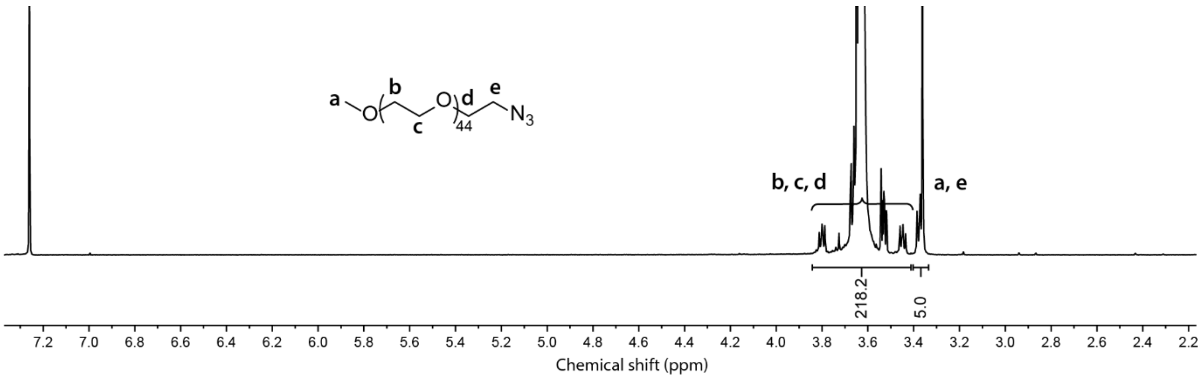

**Figure S2.** $^{1}$H-NMR spectrum of PEG-$N_3$.

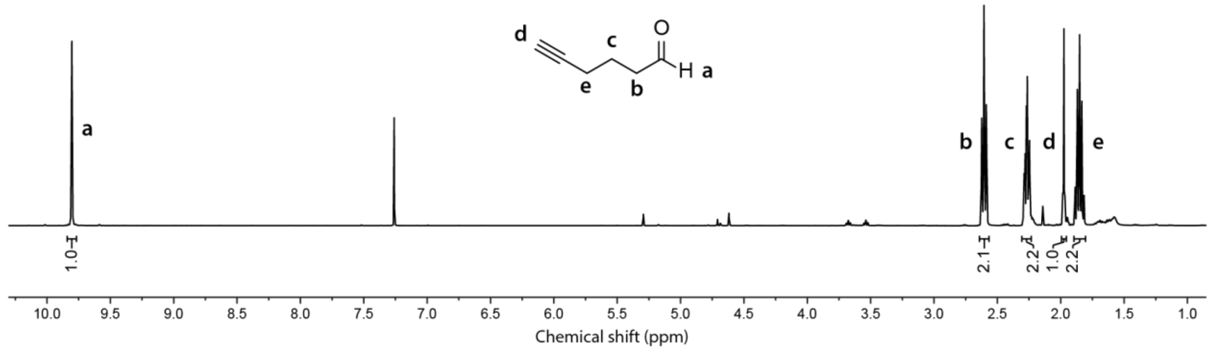

**Figure S3.** $^{1}$H-NMR spectrum of Compound **1**.

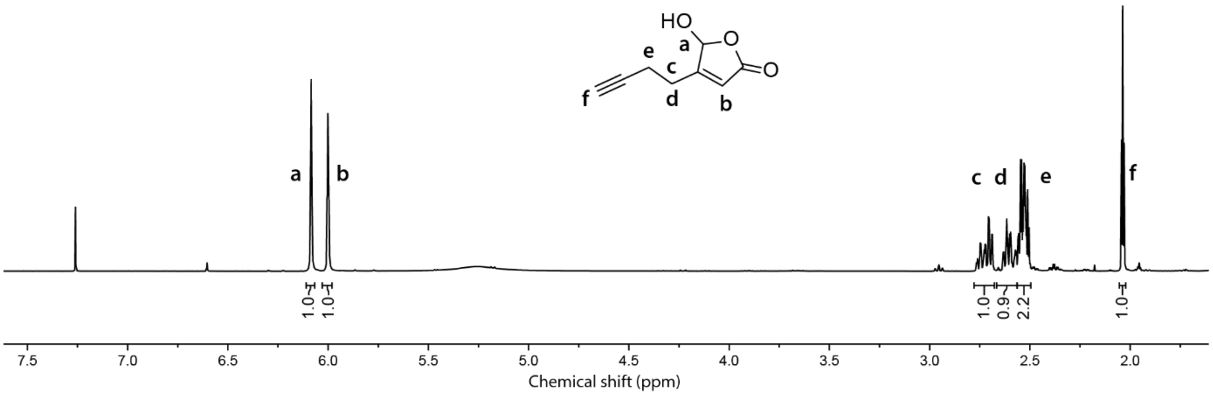

**Figure S4.** $^{1}$H-NMR spectrum of Compound **2**.

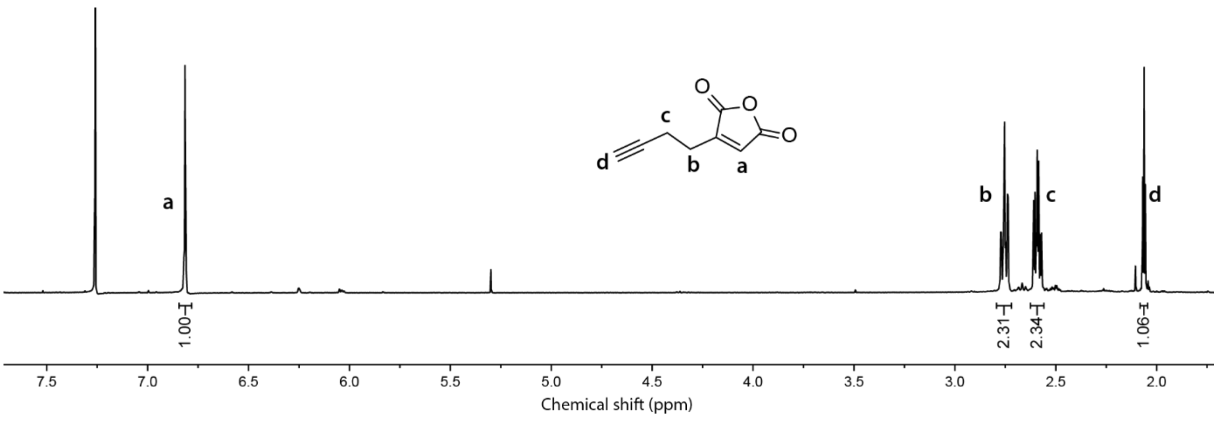

**Figure S5.** $^{1}$H-NMR spectrum of Compound **3**.

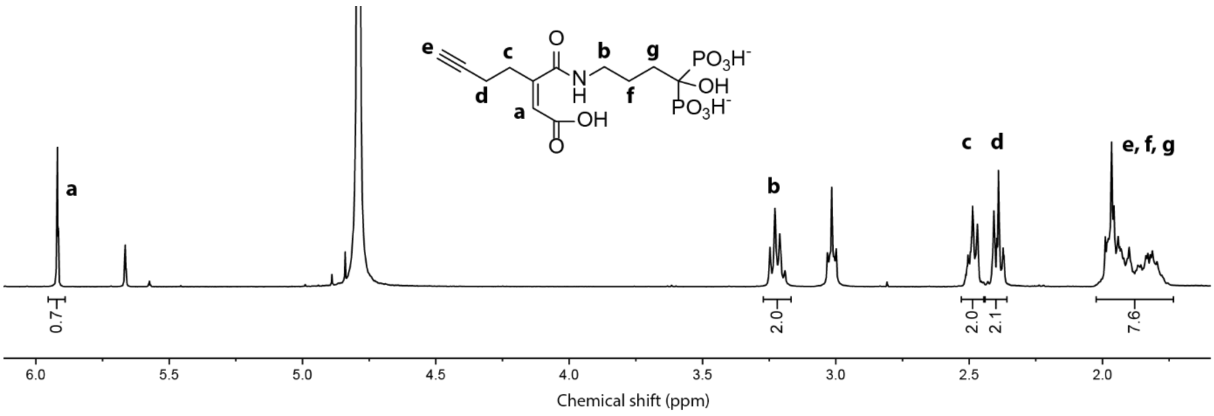

**Figure S6.** $^{1}$H-NMR spectrum of Compound **4**.

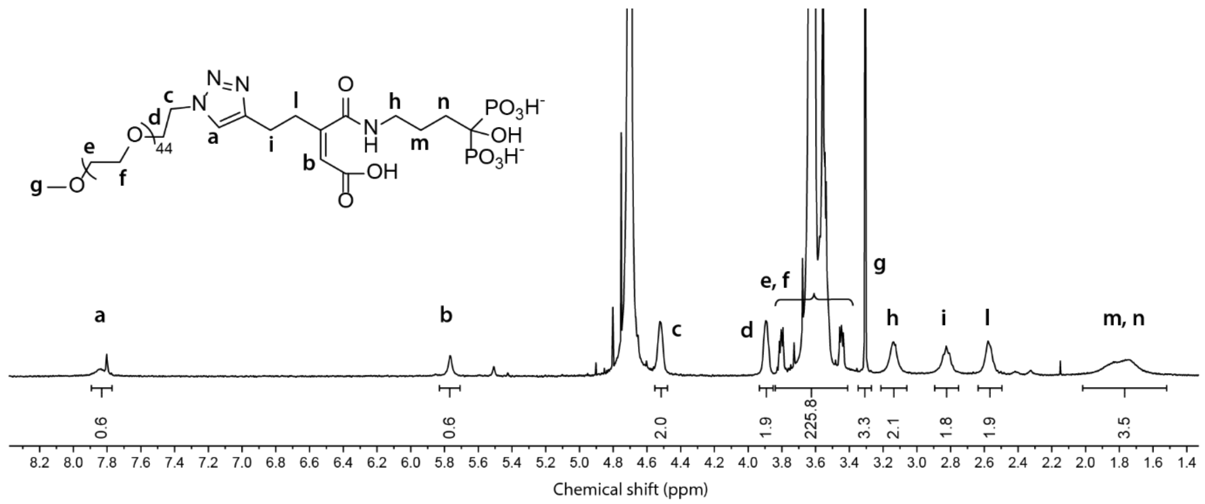

**Figure S7.** [1]H-NMR spectrum of PEG-ma-Ale.

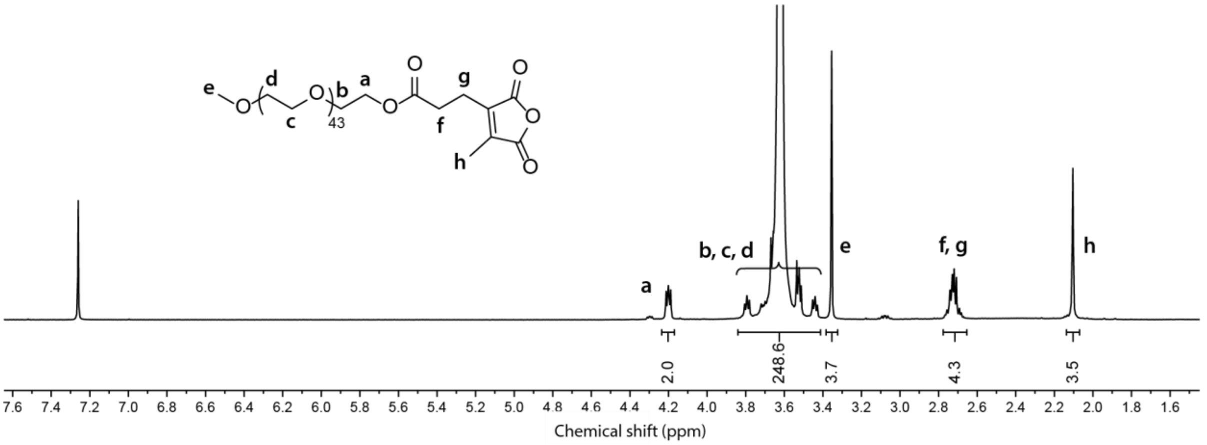

**Figure S8.** [1]H-NMR spectrum of Compound **5**.

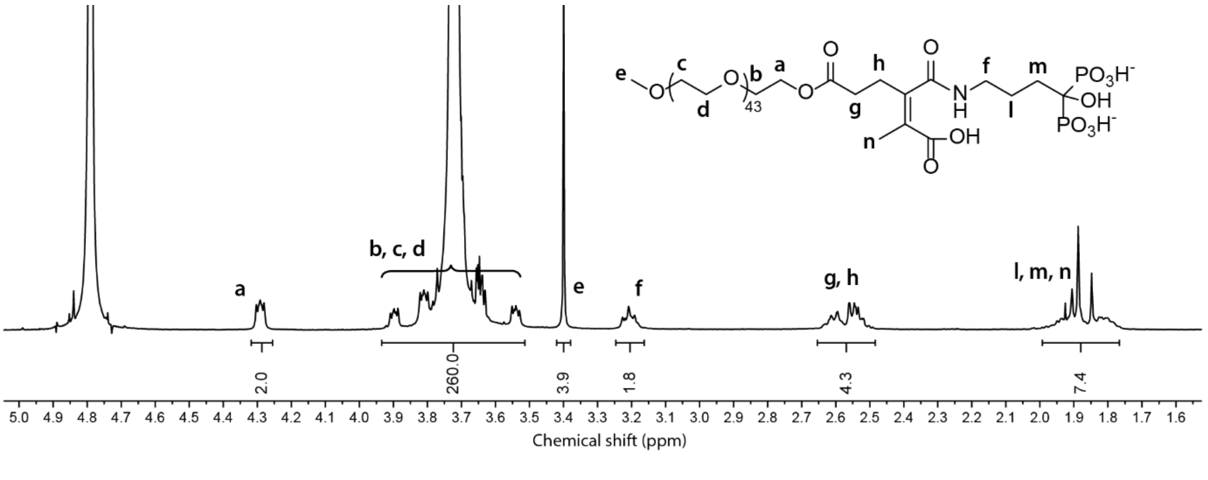

**Figure S9.** [1]H-NMR spectrum of PEG-mma-Ale.

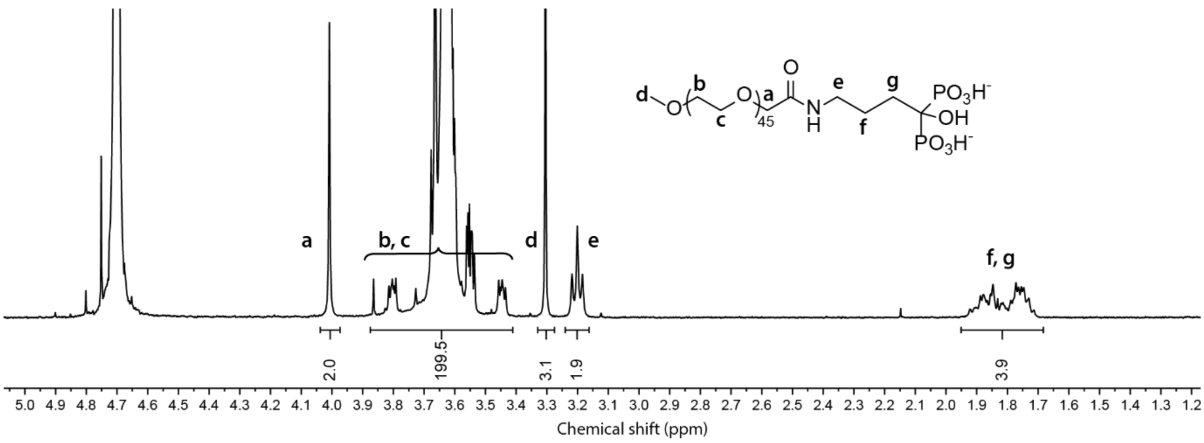

**Figure S10.** $^{1}$H-NMR spectrum of PEG-a-Ale.

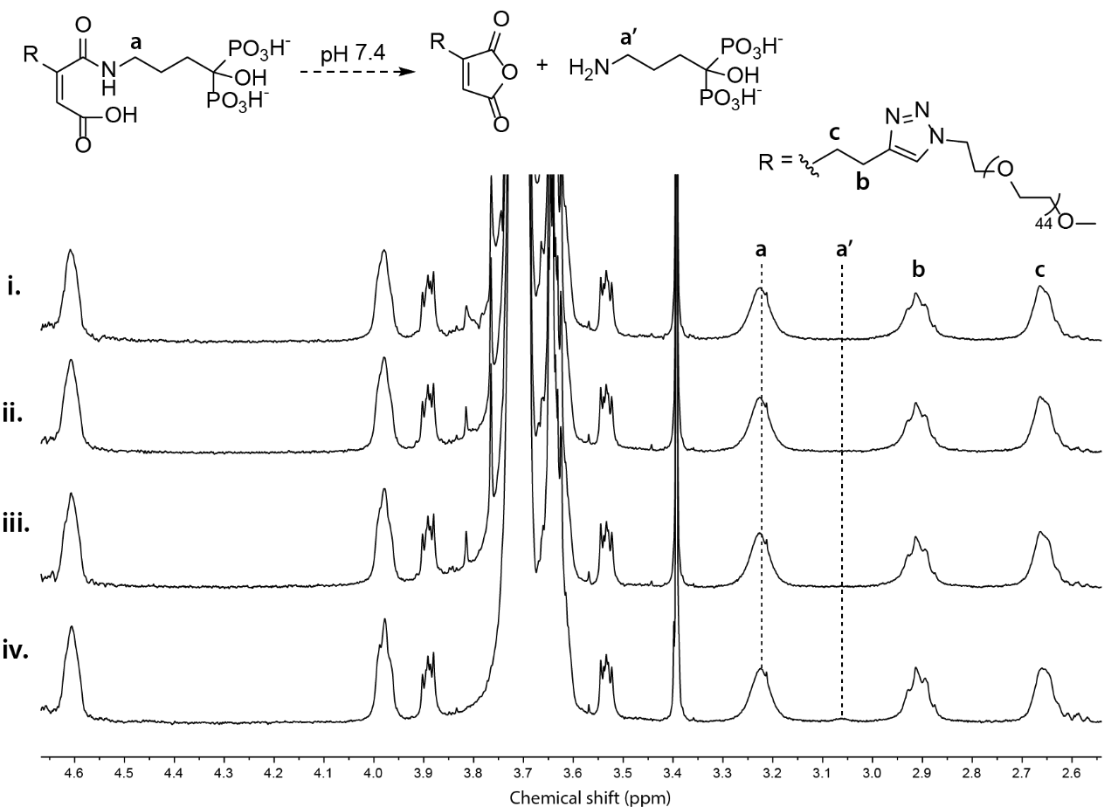

**Figure S11.** PEG-ma-Ale was stable upon incubation at pH 7.4. $^1$H-NMR spectrum of PEG-ma-Ale after 10 min incubation at pH 7.4 (i) and after 2 h (ii), 4 h (iii) or 24 h (iv) incubation at the same pH.

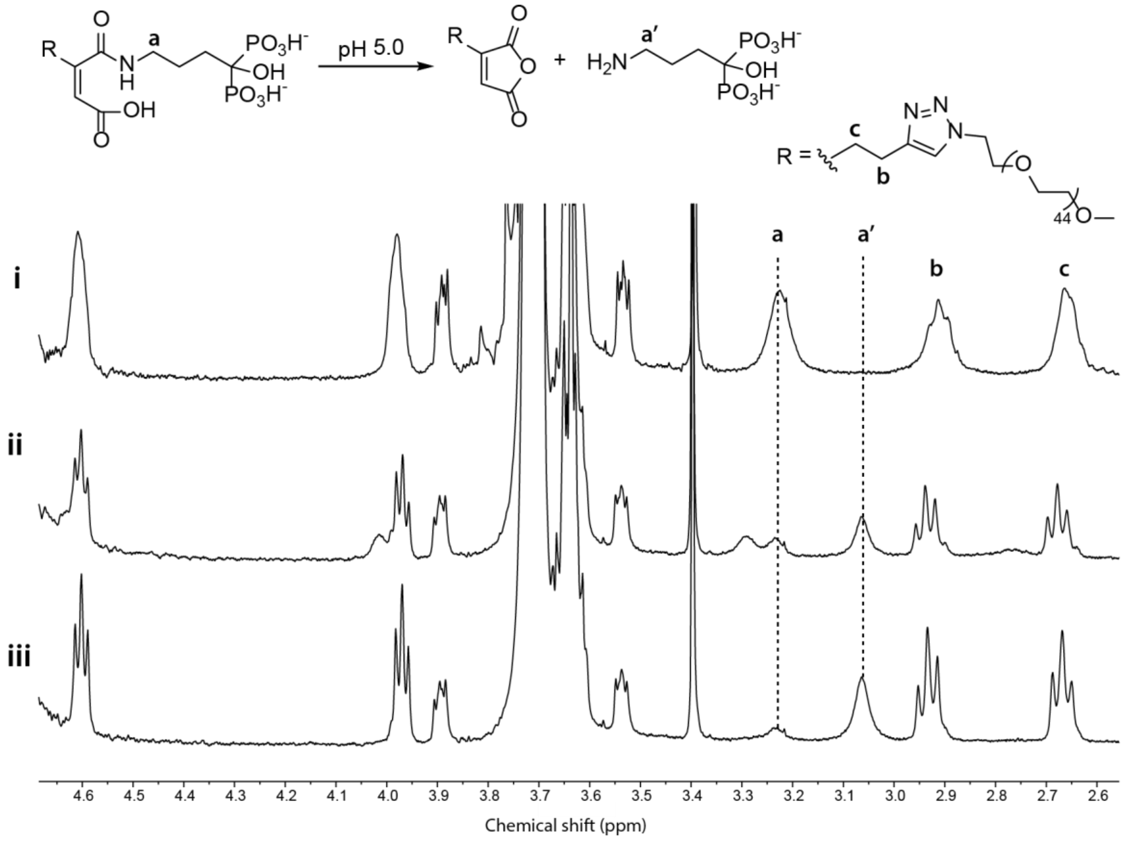

**Figure S12.** PEG-ma-Ale slowly hydrolyzed at pH 5.0. $^1$H-NMR spectrum of PEG-ma-Ale after 10 min incubation at pH 7.4 (i) and after 4 h (ii) and 24 h (iii) incubation at pH 5.0. In acidic environment, the integral of the methylene protons next to the nitrogen in the intact amide (*i.e.* 3.24 ppm) decreases, while the integral of the methylene protons close to the nitrogen in Ale (*i.e.* 3.05 ppm) increases, indicating degradation of the amide.

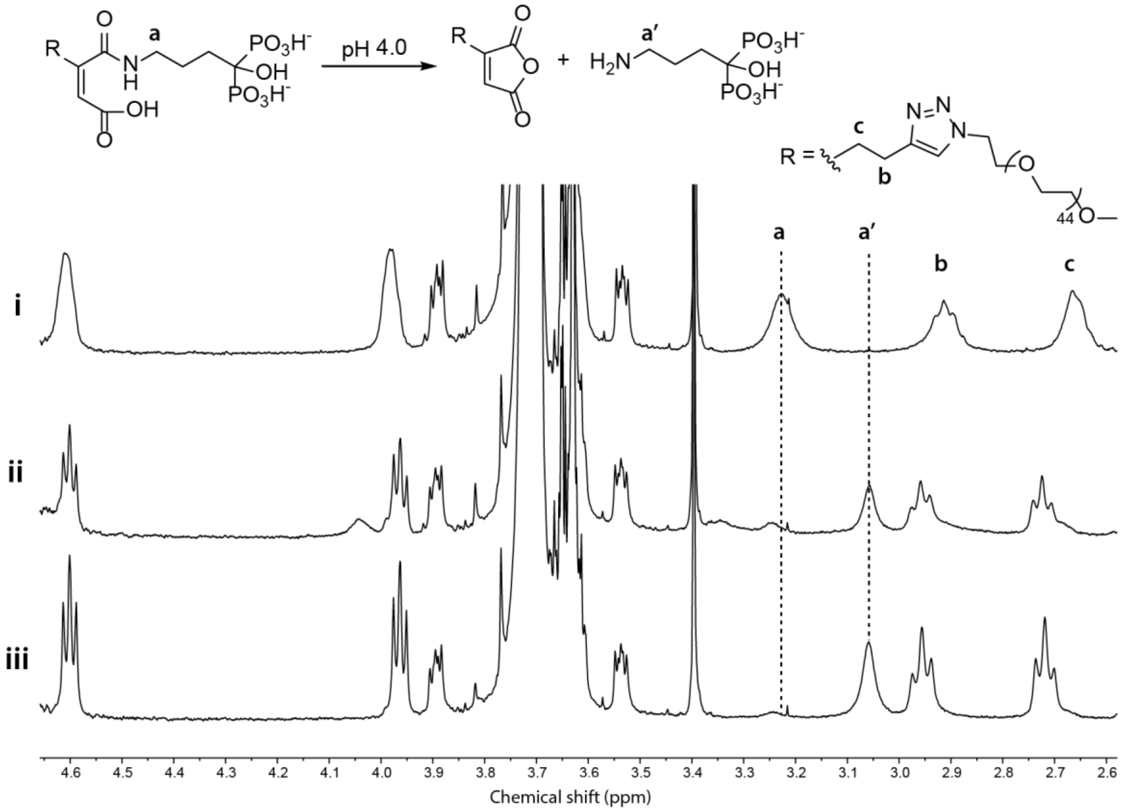

**Figure S13.** PEG-ma-Ale hydrolyzed quickly at pH 4.0. $^1$H-NMR spectrum of PEG-ma-Ale after 10 min incubation at pH 7.4 (i) and its $^1$H-NMR spectrum after 1 h incubation at pH 4.0 (ii) and 6 h incubation at pH 4.0 (iii). After 6 h, the NMR spectrum presents a prominent peak at 3.05 ppm, corresponding to the methylene protons close to the nitrogen in Ale, which is released from the conjugate.

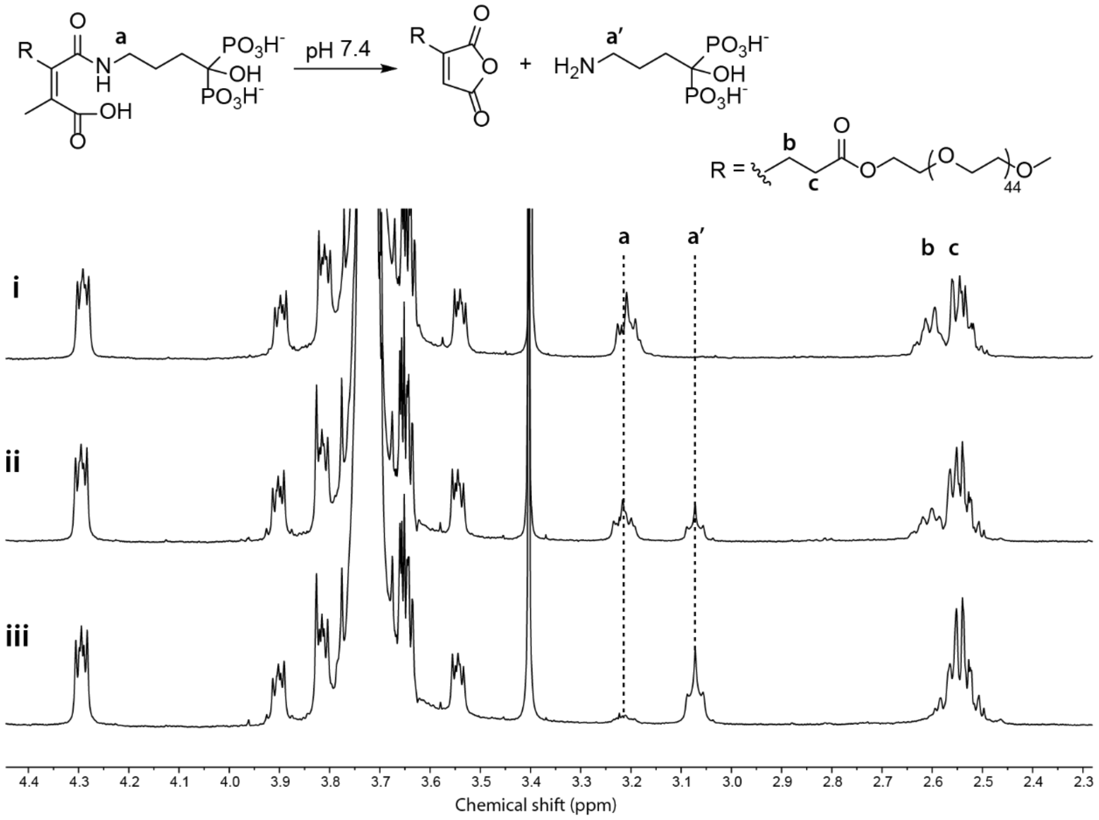

**Figure S14.** PEG-mma-Ale was quickly hydrolyzed at pH 7.4. [1]H-NMR spectrum of PEG-ma-Ale after 10 min incubation at pH 10 (i) and after 1 h (ii) and 4 h (iii) incubation at pH 7.4. The appearance of the peak at 3.05 ppm, which corresponds to the methylene protons close to the nitrogen in Ale, indicates degradation of the conjugate.

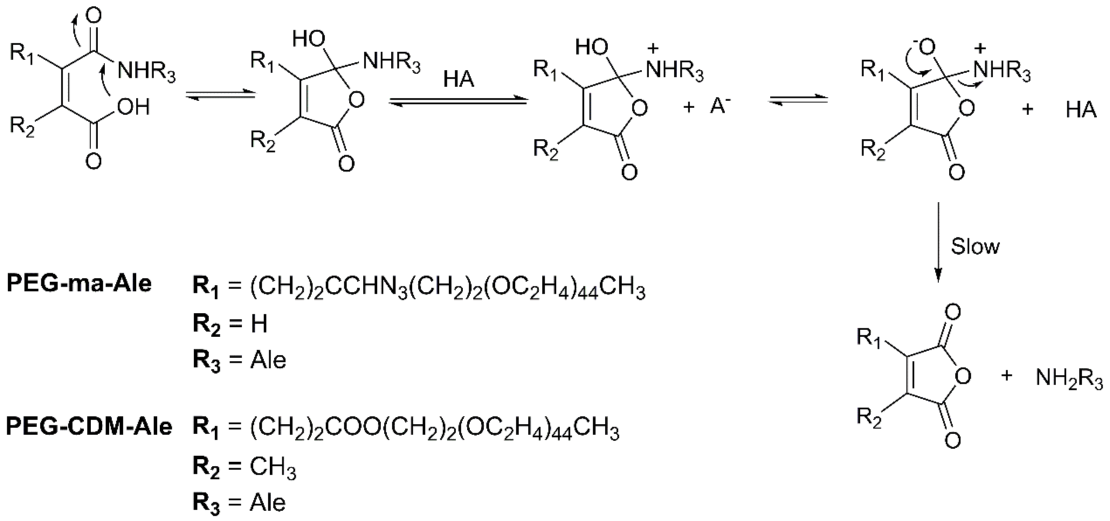

**Figure S15.** A possible degradation mechanism of PEG-ma-Ale and PEG-mma-Ale maleic acid amide derivatives in acidic conditions proposed by Kang et al [6].

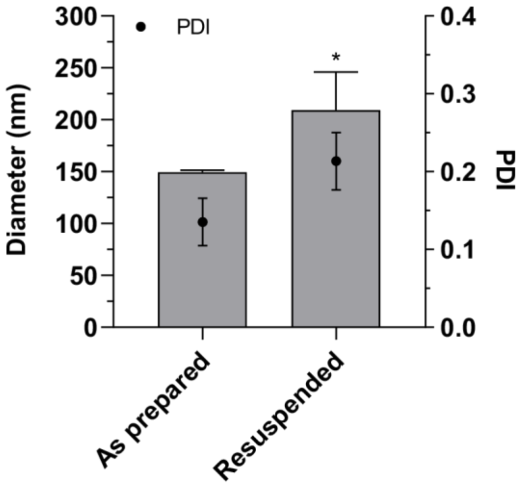

**Figure S16.** Hydrodynamic diameter of pEGFP loaded CaP NPs stabilized with 15 μM PEG-ma-Ale prepared in "low osmolality" conditions, generated using 7 mM HEPES buffer (containing 18 mM NaCl and 1.5 mM $Na_2HPO_3$, pH 7.4), 125 mM $CaCl_2$, and 2 mM TRIS, pH 7.4. After resuspension following lyophilisation, the size of the particles increased from ~150 to ~200 nm. Data are presented as mean ± SD (n=3). * $p \leq 0.05$ (refers to diameter).

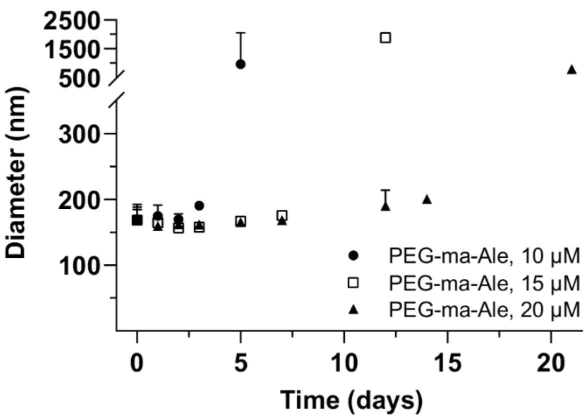

**Figure S17.** Long-term stability of PEG-ma-Ale coated CaP NPs at RT. At 10 µM PEG-ma-Ale initial concentration, CaP NPs remained stable for up to 3 days. PEG-ma-Ale concentrations of 15 and 20 µM could surface stabilize the particles for up to 7 and 15 days, respectively. Data are presented as mean + SD (n=3).

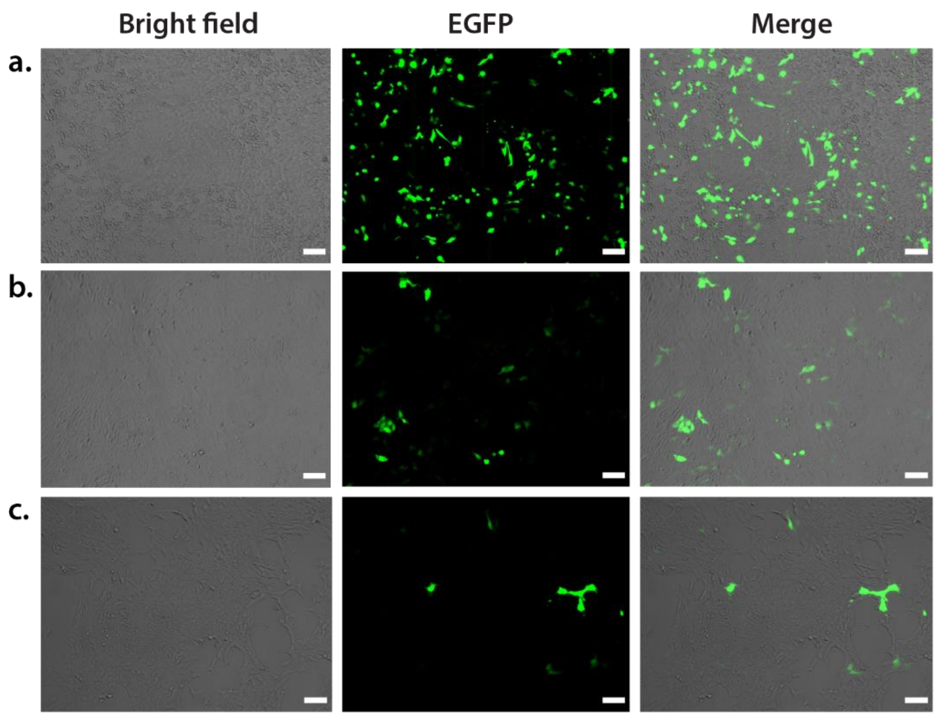

**Figure S18.** Representative fluorescent images of transfection with pEGFP: X-tremGENE™ (a), CaP NPs stabilized with 10 (b) and 20 µM (c) PEG-ma-Ale. 4T1 cells were incubated with the transfection agent for 48 h in the presence of 10% FBS. Scale bar represents 100 µm.

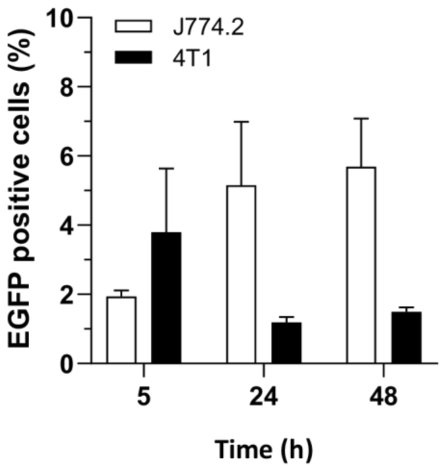

**Figure S19.** Transfection efficiency of uncoated CaP NPs after 5, 24, or 48 h incubation with J774.2 or 4T1 cells in Opti-MEM supplemented with 10% FBS. Data are presented as mean + SD (n=3).

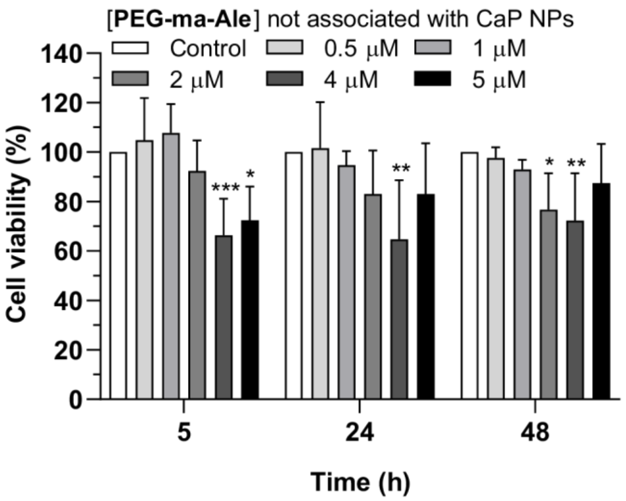

**Figure S20.** Viability of J774.2 cells after exposure to free PEG-ma-Ale for 5, 24 and 48 h. Data are presented as mean + SD (n=6-9). * $p \leq 0.05$; ** $p \leq 0.01$; *** $p \leq 0.001$ *vs.* control wells (not treated).

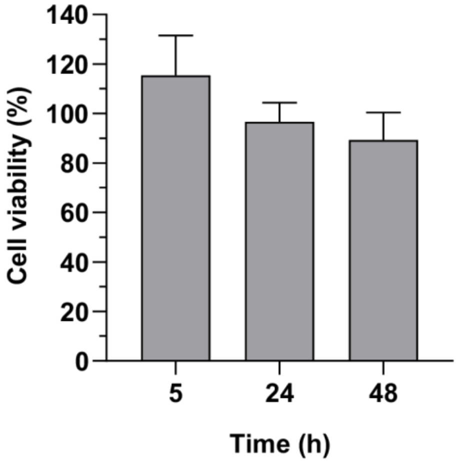

**Figure S21.** Uncoated CaP precipitates are not toxic for macrophages. Cell viability of J774.2 cells incubated for 5, 24 or 48 h with not PEGylated CaP NPs. Data are expressed as mean + SD (n=6). Difference between treated wells and control wells (not treated, cell viability = 100 %) were not statistically significant ($p > 0.05$).

**Table S1**. Physicochemical properties of PEG-a-Ale coated CaP NPs. Means ± SD (n = 3).

| | Blank CaP NPs | |
|---|---|---|
| **PEG-a-Ale (μM)** | **10** | **20** |
| **Diameter (nm)** | 139 ± 3 | 140 ± 1 |
| **PDI** | 0.11 | 0.12 |
| **ζ –potential** | -1 ± 1 | 0 ± 1 |

### S.3 Reference of supporting information
